# Topographic Axes of Wiring Space Converge to Genetic Topography in Shaping Human Cortical Layout

**DOI:** 10.1101/2023.09.06.556618

**Authors:** Deying Li, Yufan Wang, Liang Ma, Yaping Wang, Luqi Cheng, Yinan Liu, Weiyang Shi, Yuheng Lu, Haiyan Wang, Chaohong Gao, Camilla T. Erichsen, Yu Zhang, Zhengyi Yang, Simon B Eickhoff, Chi-Hua Chen, Tianzi Jiang, Congying Chu, Lingzhong Fan

## Abstract

Genetic information is involved in the gradual emergence of cortical areas since the neural tube begins to form, shaping the heterogeneous functions of neural circuits in the human brain. Informed by invasive tract-tracing measurements, the cortex exhibits marked interareal variation in connectivity profiles to reveal the heterogeneity across cortical areas. However, it remains unclear about the organizing principles possibly shared by genetics and cortical wiring to manifest the spatial heterogeneity across the cortex. Instead of considering a complex one-to-one mapping between genetic coding and interareal connectivity, we hypothesized the existence of a more efficient way that the organizing principles are embedded in genetic profiles to underpin the cortical wiring space. Leveraging on the vertex-wise tractography in diffusion-weighted MRI, we derived the global connectopies to reliably index the organizing principles of interareal connectivity variation in a low-dimensional space, which specifically captured three dominant topographic patterns along the dorsoventral, rostrocaudal, and mediolateral axes of the cortex. More importantly, we demonstrated that the global connectopies converge to the gradients of vertex-by-vertex genetic correlation matrix on the phenotype of cortical morphology and the cortex-wide spatiomolecular gradients. By diving into the genetic profiles, we found the critical role of genes scaffolding the global connectopies were related to brain morphogenesis and enriched in radial glial cells before birth and excitatory neurons after birth. Taken together, our findings demonstrated the existence of a genetically determined space to encode the interareal connectivity variation, which may give new insights into the links between cortical connections and arealization.

## Introduction

Mounting evidence has suggested that the genetic effects on the anatomical phenotypes of the human brain are spatially heterogeneous ^1–4^, demonstrating the vital roles of genes in establishing the configuration of brain space, such as the anatomical hierarchy ^5,6^ and the wiring diagram ^7–9^. The profile of cortical wiring that can be characterized by using neuroimaging-based tractography is especially reliable for indicating the heterogeneity across cortical areas ^10,11^, reflecting the differentiation in local microstructures and connectivity patterns. The interareal connectivity variation has further revealed the spatially heterogeneous patterns of cortical organization, such as the regional controllability related to cognitive dynamics ^12,13^. In contrast, the deficit of interareal connectivity has demonstrated the specificity of the behavioral manifestations of the brain lesion ^14,15^. Although spatial heterogeneity has been separately found to be embedded in genetic architecture and interareal connectivity variation, it is still largely unknown if a unified organizing principle exists underlying their indication of spatial heterogeneity.

The spatiotemporal distribution of genetic factors along the developing brain is found to set the primary cues to guide the process of cortical arealization ^16,17^. It has been pointed out that gene-coded signalling molecules and transcriptional profiles may contribute to this guidance process, and these mechanisms may even be conserved into maturity ^18,19^. Specifically, under the effect of morphogens secreted in the patterning centers, transcription factors are expressed in a graded manner and determine the areal fate and the expression of cell-surface molecules, thus determining the topographic organization of synaptic inputs and outputs ^1,20^ and directing axonal outgrowth ^21^. Nevertheless, considering the complexity and flexibility of neural circuits, it would be difficult to conceive the existence of a one-to-one mapping between the genetic codes and the wiring patterns^22^. Alternatively, we hypothesize that the genetic profiles distributed across the cortex may contain the organizing principles of cortical wiring ^22^. In this way, the genetic processes and the cortical wiring patterns can be systematically unified to provide a more complete understanding of the heterogeneity across the cortex. Recent advances in acquiring high-throughput transcriptomic data of the human brain ^23^ and large-scale twin samples ^2,3^ with neuroimaging data provide essential resources to test our hypothesis.

Furthermore, even if we can identify the genetic topography, it remains unknown whether we can leverage it to explain the interareal connectivity variation captured by high-resolution connectomics ^24,25^. Recent studies revealed that cortical areas are hierarchically along the cortex from a large-scale gradient perspective of connectivity ^26,27^, indicating that the cortex is organized topologically within interareal connectivity variation, supporting diverse dynamics and functions ^28–31^. However, traditional neural circuit tracing techniques, including classical tracers or virus tracing, are precluded in humans, thereby creating the need for noninvasive measures of the topography embedded in interareal connectivity variation, which would contribute to testing the existence of unified rules underlying the genetic topography and interareal connectivity variation.

To address these open questions, we first quantified the interareal connectivity variation by establishing the similarity matrix of the structural connectivity profiles. Leveraging a manifold learning approach, we identified three components (hereafter also referred to as global connectopy) running separately along the dorsoventral, rostrocaudal, and mediolateral axes of the brain anatomy, which efficiently delineated the interareal connectivity variation. Then, we characterized the genetic topography by respectively utilizing the gradients from the twin-based genetic correlation matrix of cortical morphology and the cortex-wide gene expression matrix. Based on the identified global connectopy and the genetic topography, we provided evidence supporting our hypothesis by demonstrating the significant consistency between them. Furthermore, we identified specific genes associated with the global connectopies which were involved with brain morphogenesis and enriched in radial glial cells before birth and excitatory neurons related with cortical projection circuit formation after birth. Our analytic logic is detailed in Figure 1 to overview the current study.

**Figure 1.**
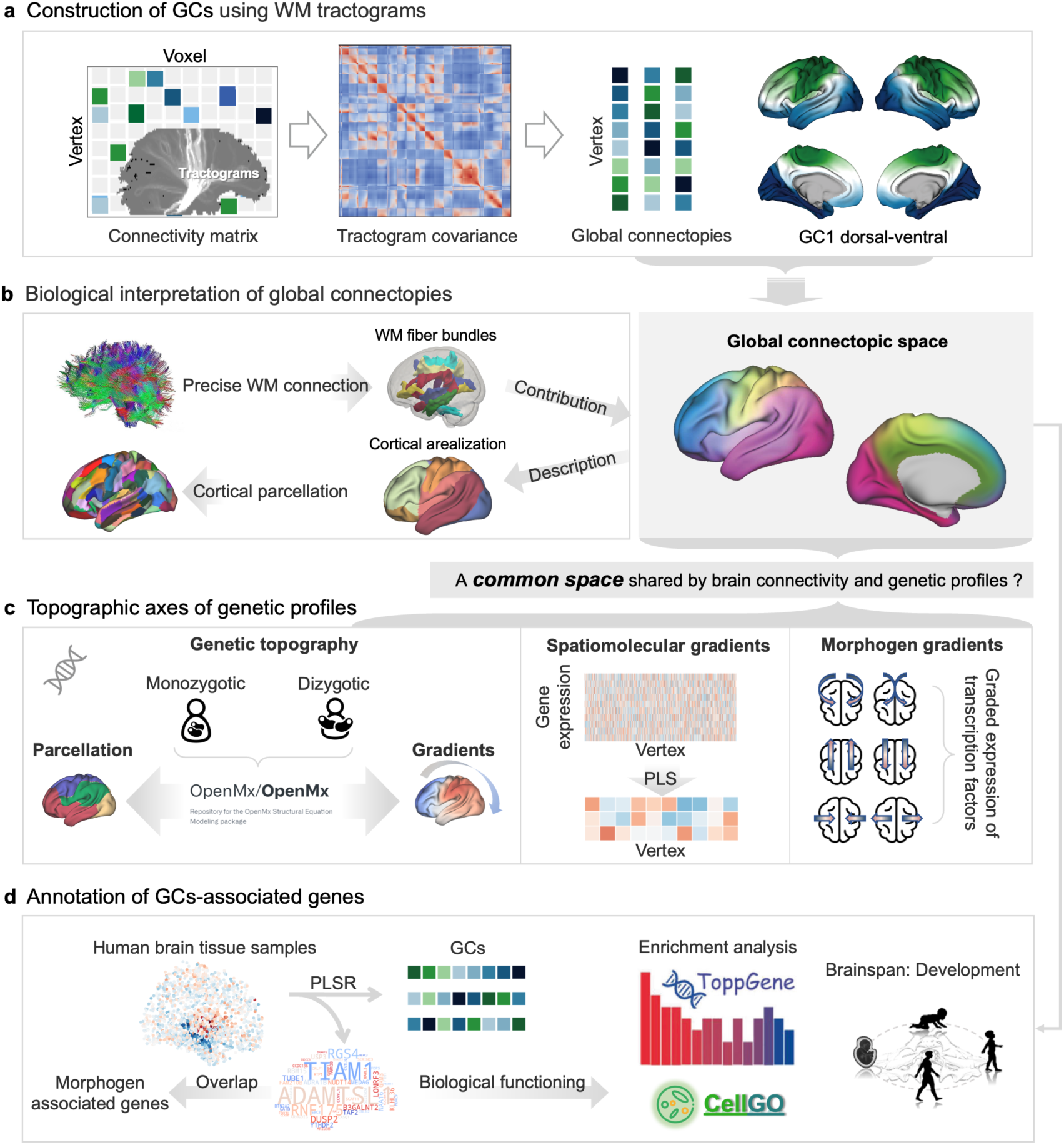
Overview of the analysis pipeline. We investigated the hypothesis that the existence of a common space shared by brain connectivity and genetic profiles. **(a)** The construction of global connectopies (GCs). White matter tractograms were calculated to get a similarity matrix, i.e., a tractogram covariance (TC) matrix. Diffusion embedding was implemented on the TC matrix, resulting in low-dimensional gradients. **(b)** Biological interpretation of the GCs. White matter bundles that contribute to the GCs were analyzed (top panel). Global connectopies were demonstrated to provide a large-scale descriptor of cortical cartography, which may give insight into cortical parcellation ^10^. **(c)** Topographic axes of genetic profiles. We demonstrated a correspondence between the genetic influence on cortical morphology and global connectopies and established that the three global connectopies are consistent with morphogen gradients in the developing brain and spatiomolecular gradients in adulthood and provided evidence that specific genes drive the formation of the cortical organization. **(d)** Annotation of GCs-associated genes. Genes associated with the GCs were identified and input to enrichment and development analyses.

## Results

### Three global connectopies in the human brain

To explore how the spatial organization of the human brain is shaped by the underlying structural connections with the white matter, we characterized the global connectopies across the whole brain. We first built a vertex-wise similarity matrix of structural connectivity using 100 unrelated subjects from the Human Connectome Project dataset ^32^. Specifically, structural connectivity profiles were obtained from probabilistic tractography along ∼30k vertices for each hemisphere and were correlated between each pair of vertices. Tractogram covariance (TC) matrices were thresholded at 0 and averaged across subjects. In short, the TC matrix captured structural connectivity similarity across the whole brain.

We implemented diffusion embedding on the TC matrix, a manifold learning method previously used to capture functional gradients ^33^. The resultant components revealed the position of vertices along the axes that had the most dominant differences in the given structural connectivity profile. The first three global connectopies separately showed dorsoventral, rostrocaudal, and mediolateral patterns and together accounted for 33% of the total variance (Figure 2a).

**Figure 2.**
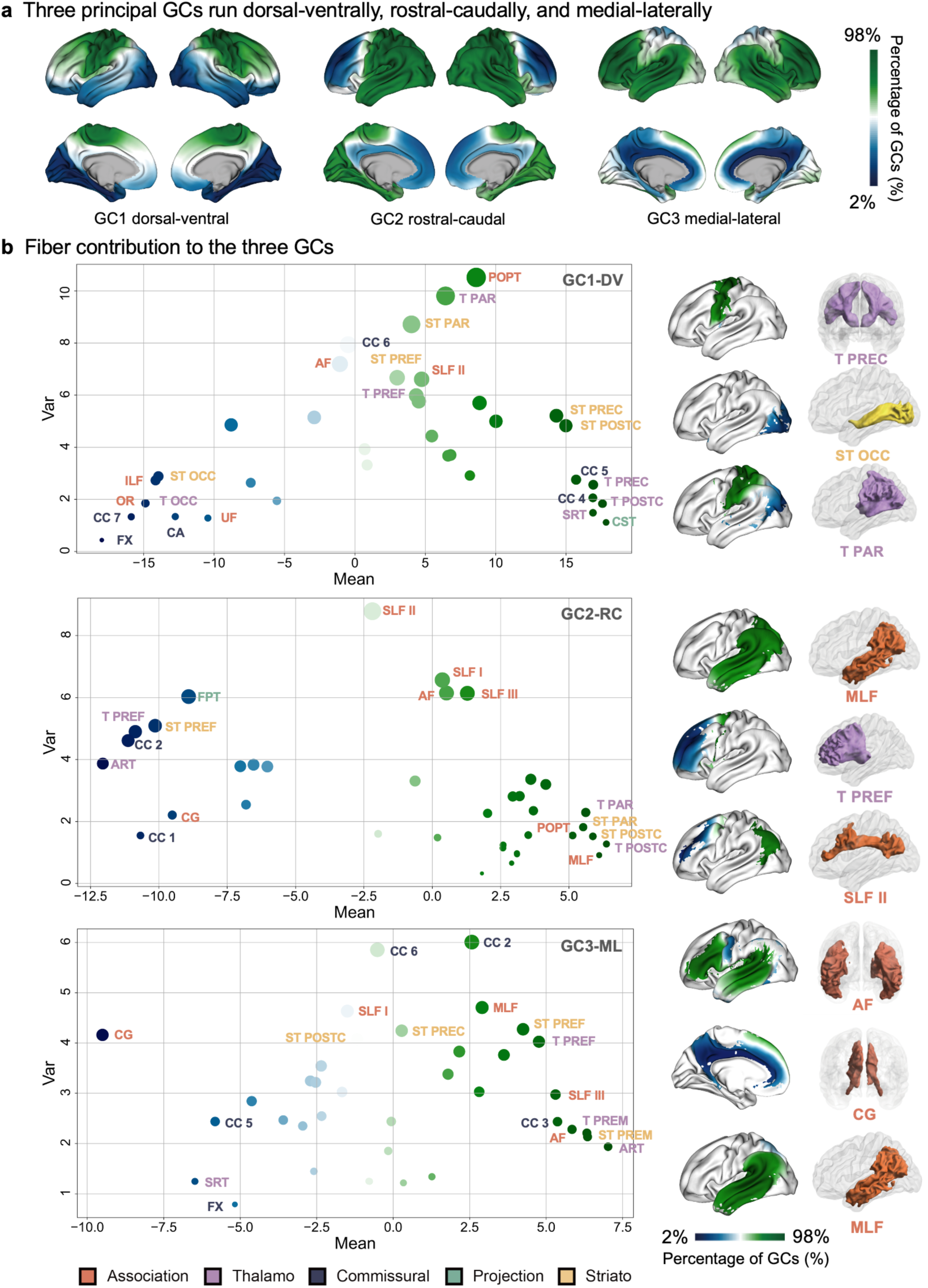
The three global connectopies and white matter tracts contribute to the GCs. **(a)** The first three global connectopies ran dorsoventrally, rostrocaudally, and mediolaterally and were termed GC1-DV, GC2-RC, and GC3-ML, respectively. **(b)** The three global connectopies are situated by distinct sets of white matter tracts. Mean values and variances of the contribution of five types of tracts to the global connectopies are shown. Each dot represents one tract. Different colors represented different types of tracts, with deeper colors for higher contributions. The right two columns show two types of characteristic tracts contributing to the global connectopies, with one type of tract situated at one extreme of the global connectopy and the other stretched across the global connectopy. The first column shows the cortical projection on the surface of the tract, and the second shows the corresponding fiber bundles reconstructed by TractSeg.

More specifically, the first global connectopy (Global connectopy 1 dorsoventral, GC1-DV), following a dorsoventral pattern, was anchored on one end by the occipital, inferior temporal, and orbital frontal cortex and on the other end by the motor and sensorimotor cortex. The second global connectopy (Global connectopy 2 rostrocaudal, GC2-RC) varied along a rostrocaudal axis, radiating from the occipitoparietal cortex and ending in the prefrontal cortex. The third global connectopy (Global connectopy 3 mediolateral, GC3-ML) showed a mediolateral pattern, with the highest expression in the lateral temporal and prefrontal cortex and the lowest in the cingulate cortex (Figure 2a).

These patterns were stable regardless of the number of subjects considered (Figure S1) and were not affected by age, sex, or brain size (Figure S2). Individual GCs were computed for each subject and aligned to the group GC with Procrustes rotation, allowing for comparison between individuals. The individual global connectopies highly correlated with the group GC (all *r* > 0.9; Figure S3). We further showed that global connectopies are beyond geodesic distance (Figure S4) and demonstrated the role of long-range connections in the formation of global connectopies (Figure S5).

### The three connectopies are situated by distinct sets of white matter tracts

We further explored how the underlying white matter shaped these spatial organizations. We fitted general linear models (GLMs) to interpolate measures from the gray matter into the white matter ^34^. Specifically, we used the global connectopies as a dependent variable and the structural connection matrix as an independent variable, thus projecting each global connectopy onto the white matter (Figure S6). For GC1-DV, the white matter voxels were distributed along the ventral-dorsal axis beneath the surface. Specifically, voxels lying near the sensorimotor/motor cortex and occipitotemporal cortex showed values that were markedly consistent with the locations of the extremes of the first global connectopy. Voxels corresponding to GC2-RC and GC3-ML varied from posterior to anterior and medial to lateral.

We then focused on the specific tracts related to the emergence of the global connectopies. We reconstructed 72 fiber bundles following TractSeg ^35^ (Table S1) and built group-level connectivity blueprints ^36^, Where the rows revealed the cortical termination patterns of the tracts. We first extracted the projection on the surface of each tract and calculated the mean value and variance of connectopy values of tract projection on the surface (Figure 2b). We identified the tracts that were either situated at one of the extremes of the global connectopy or were spread out along the axis. The results for the bilateral tracts were averaged between the two hemispheres and are shown in the scatterplot (Figure 2b; detailed values are shown in Figure S7).

Two types of characteristic tracts are shown in the right columns of Figure 2b, including thalamo-precentral tract (T_PREC), striato-occipital tract (ST_OCC), and thalamo-parietal tract (T_PAR) for GC1-DV as well as middle longitudinal fascicle (MLF), thalamo-prefrontal (T_PREF), superior longitudinal fascicle II (SLF_II) for GC2-RC along with arcuate fascicle (AF), cingulum (CG), and MLF for GC3-ML.

For GC1-DV, T_PREC, which connects the thalamus with the precentral gyrus, is related to motor task performance ^37^. T_PREC has a distinctly vertical expanding shape and showed high mean values near the dorsal end of GC1-DV, thus expressing the most significant contribution to the global connectopy. ST_OCC connects the striatum with the occipital cortex, showing high mean values at the ventral end and having a “stretching” impact on the global connectopy pattern. In contrast, T_PAR has a high variance within the GC1-DV and appears to constrain the global connectopy formation.

For GC2-FC, the second branch of the superior longitudinal fascicle, which mostly runs from the posterior to the anterior, starting from the middle frontal gyrus and terminating in the angular gyrus, had the greatest mean value for the white matter tract. The SLF_II, which covered most of the areas in the second global connectopy, also showed high values, providing the prefrontal cortex with parietal cortex information about the perception of visual space and providing information about working memory in the prefrontal cortex to the parietal cortex to focus spatial attention and regulate the selection of spatial information ^38^. Other “high-contribution” tracts, including the MLF and T_PREF, are either in the anterior or posterior regions.

The GC3-ML showed a similar pattern to the tracts that made high contributions. The corpus callosum, the giant white matter fiber bundle consisting of millions of axonal projections, spans from the medial brain through the different parts of the cortex and connects all the brain lobes. Thalamo- and striato-cortical projections also project from the subcortical nuclei and reach the different lateral cortices. We suggest that the former type of tract plays a role in stretching the brain along the axis while the latter type has a constraining effect and that these work together to shape the organization of the brain.

### Global connectopies provide a large-scale descriptor of arealization

To inspect the global connectopies as a unit, we normalized three connectopy values at each vertex to generate RGB values and mapped them on the surface to make lobes or subregions seen distinctly (Figure 3a). We termed this map, the global connectopic space, which quantified the topography of the dominant global connectopy. The cortex, shown in red, green, and blue, indicated the individual regions that were dominated by each global connectopy. The regions that were dominated by a combination of more than one global connectopy were done in different colors, e.g., the “yellow” region was dominated by the first and second global connectopies together, as shown in the cube in Figure 3a.

**Figure 3.**
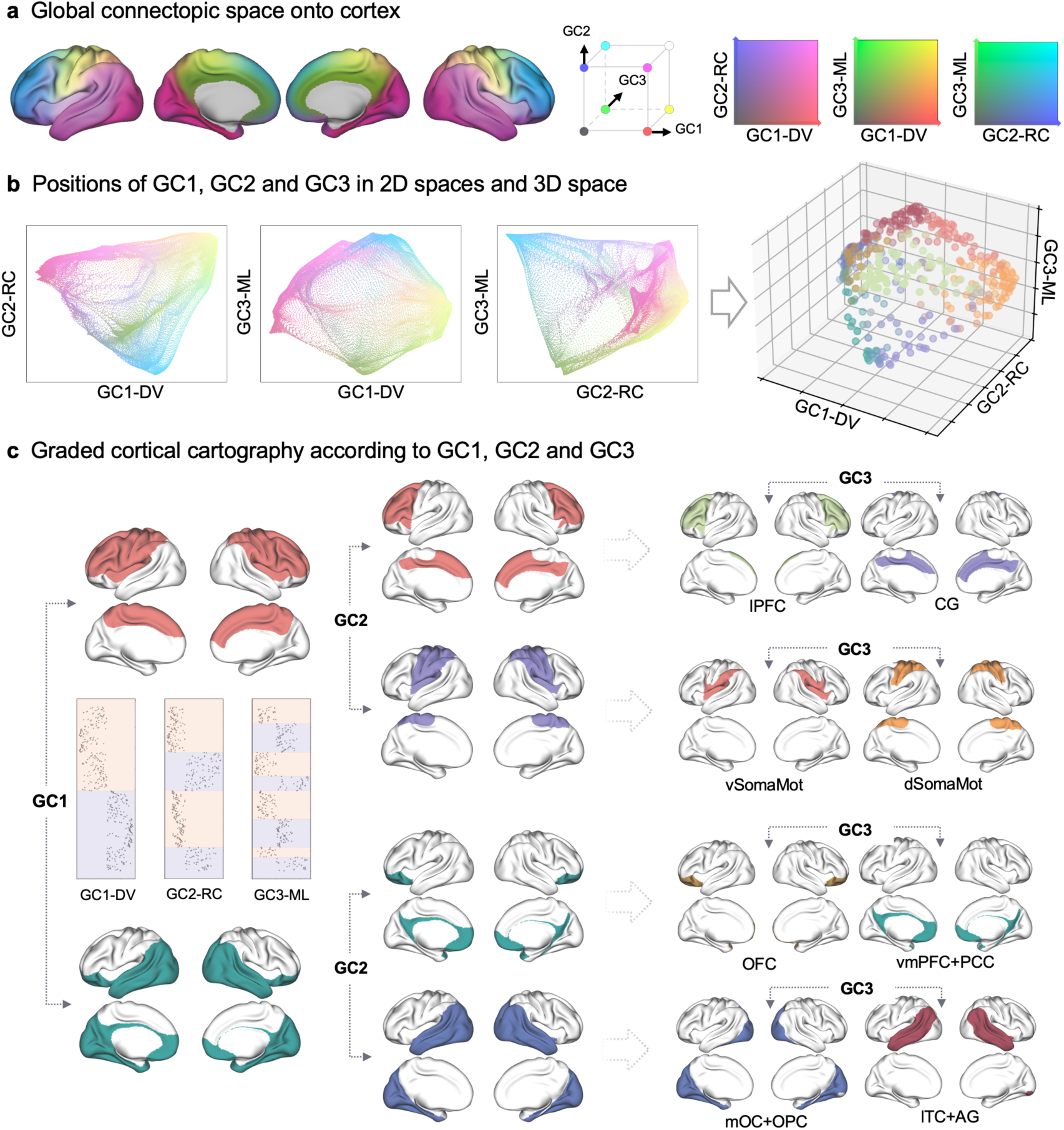
Global connectopies provide a large-scale descriptor of arealization. **(a)** The three global connectopies were plotted together in what we termed the global connectopic space, with connectopy values assigned to the RGB values at each vertex. The global connectopic space integrates the information of three global connectopies and quantifies the topography of the dominant global connectopy. **(b)** Left: Several lobes or subregions clearly appear when every two global connectopies are plotted together on the 2D space, with the extremes of GCs clearly and continuously shown. GC1-DV and GC2-RC split the cortex into the prefrontal cortex and limbic cortex, sensorimotor and motor cortex, and occipitotemporal cortex. GC1-DV and GC3-ML split the cortex into the frontal cortex, limbic cortex, and occipitotemporal cortex. GC2-RC and GC3-ML split the cortex into the prefrontal cortex, limbic cortex, and other regions. Right: The three GCs were plotted together in the 3D space. Vertices were assigned different colors according to the signs of the three axes. **(c)** Modules identified by hierarchical clustering using the three global connectopies. At each level, the brain was partitioned into two modules according to the positive and negative signs of the global connectopy. Eight modules, including lobes or functional networks, emerged clearly. vSomaMot ventral somatomotor cortex, dSomaMot dorsal somatomotor cortex, lPFC lateral prefrontal cortex, CG cingulate gyrus, lTC lateral temporal cortex, AG angular gyrus, mOC medial occipital cortex, OPC occipital polar cortex, OFC orbitofrontal cortex, vmPFC ventromedial prefrontal cortex, PCC posterior cingulate cortex.

We then plotted every pair of global connectopies in a 2D space and assigned colors based on the global connectopic space in Figure 3a. The first and the second global connectopies interacted to split the whole brain into the prefrontal cortex and limbic cortex, sensorimotor and motor cortex, and occipitotemporal cortex (Figure 3b). Similarly, the frontal cortex, limbic cortex, and occipitotemporal cortex were shown when the first and the third global connectopies were considered together, and the prefrontal cortex, limbic cortex (Figure 3b), and others were shown when the second and the third connectopy were combined (Figure 3b). The three GCs were also plotted together in the 3D space. Vertices were assigned different colors according to the signs of the three axes (Figure 3b, right).

In order to show the modules clearly in the global connectopic space, we also partitioned the brain by hierarchical clustering using the three global connectopies (Figure 3c). At the first level, the brain was partitioned into two modules based on the positive and negative signs of the first global connectopy. These two modules coincided with the division between the dorsal and ventral brain regions. Each module was further partitioned into two modules at each level by the sign of the global connectopies. The rostrocaudal and mediolateral patterns emerged after the corresponding global connectopy was considered. In the end, eight modules were captured. These were the ventral and dorsal somatomotor cortex, lateral prefrontal cortex, cingulate gyrus, lateral temporal cortex + angular gyrus, medial occipital cortex + occipital polar cortex, orbitofrontal cortex, and ventromedial prefrontal cortex + posterior cingulate cortex (Figure 3c).

### Potential genetic basis underlying global connectopies

Having established the global connectopies in the human brain, we first explored the relationship between the genetic influence on cortical morphology with the identified global connectopies. We replicated the global connectopies using twin data from the HCP and found they show high similarity with the results derived from unrelated subjects (left: *r*_G1-DV_ = 0.99, *r*_G2-RC_ = 0.99, *r*_G3-ML_ = 0.99; right: *r*_G1-DV_ = 0.99, *r*_G2-RC_ = 0.98, *r*_G3-ML_ = 0.98; Figure S8). We estimated the pairwise genetic correlation between vertices on the cortex, which revealed shared genetic influences on cortical thickness and relative area expansion between cortical vertices. The first three gradients decomposed from the genetic correlation matrix of cortical thickness showed a rostral-caudal axis, a medial-lateral axis, and a dorsal-ventral axis, respectively (left: *r*_GC1-DV, GG3-Thickness_ = 0.72, *p*_spin_ < .0028, FDR corrected; *r*_GC2-RC, GG1-Thickness_ = 0.91, *p*_spin_ < .0001, FDR corrected; *r*_GC3-ML, GG2-Thickness_ = 0.72, *p*_spin_ < .0041, FDR corrected; right: *r*_GC1-DV, GG3-Thickness_ = 0.73, *p*_spin_ < .0044, FDR corrected; *r*_GC2-RC, GG1-Thickness_ = 0.88, *p*_spin_ < .0098, FDR corrected; *r*_GC3-ML, GG2-Thickness_ = 0.71, *p*_spin_ < .1566, FDR corrected; Figure 4a, 4b). The first three gradients decomposed from the genetic correlation matrix of the surface area correspond with the three global connectopies (left: *r*_GC1-DV, GG1-Area_ = 0.53, *p*_spin_ < .1630; *r*_GC2-RC, GG2-Area_ = 0.49, *p*_spin_ < .0006, FDR corrected; *r*_GC3-ML, GG3-Area_ = 0.51, *p*_spin_ < .1031; right: *r*_GC1-DV, GG1-Area_ = 0.58, *p*_spin_ < .2131; *r*_GC2-RC, GG2-Area_ = 0.46, *p*_spin_ < .0177; *r*_GC3-ML, GG3-Area_ = 0.51, *p*_spin_ < .0812; Figure S9). Each parcel derived from the fuzzy clustering of the genetic correlation matrix of cortical thickness ^2^ also showed significant overlap with one of the four parcels derived from the hierarchical clustering identified using the global connectopies (Figure 4c, Figure S10a, b and Figure S11a, b). The overlap between parcels derived from the hierarchical clustering and parcels from the genetic correlation of surface area is additionally shown in Figure S10c, d and Figure S11c, d. These significant correspondence levels indicated a close genetic relatedness of the global connectopies.

**Figure 4.**
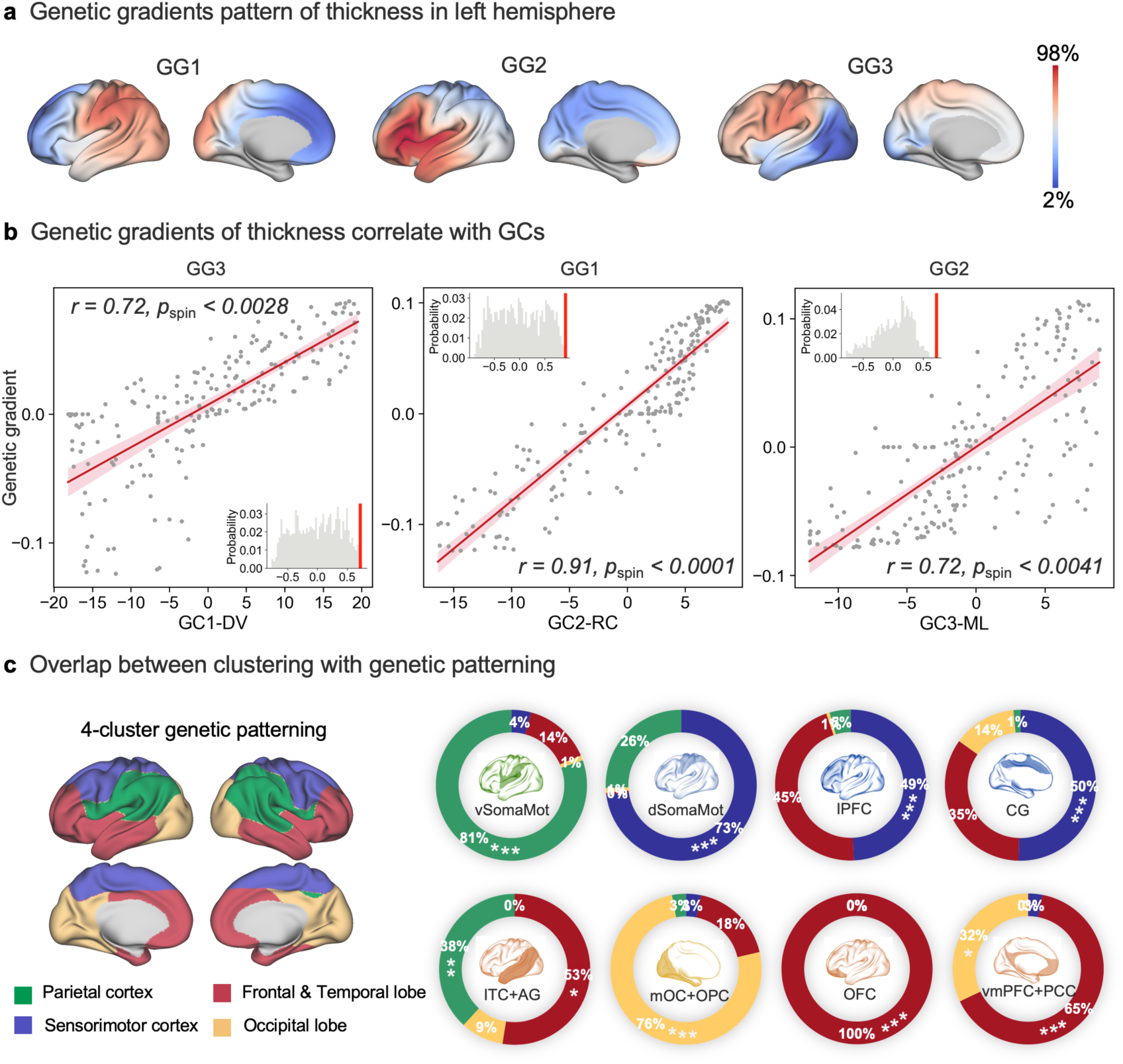
Global connectopies correspond to genetic topography. **(a)** The first three gradients of genetic similarity of cortical thickness. The genetic similarity matrix was calculated by fitting bivariate ACE models to compute the genetic correlations of the cortical thickness between two vertices in a twin dataset. **(b)** The three gradients of genetic similarity of cortical thickness show a high correlation with global connectopies (*r*_GC1-DV, GG3-Thickness_ = 0.72, *p*_spin_ < .0028, FDR corrected; *r*_GC2-RC, GG1-Thickness_ = 0.91, *p*_spin_ < .0001, FDR corrected; *r*_GC3-ML, GG2-Thickness_ = 0.72, *p*_spin_ < .0041, FDR corrected). **(c)** Overlap between four modules derived from hierarchical clustering with genetic patterning of cortical thickness. Four-cluster genetic patterning was obtained by performing fuzzy clustering on the genetic similarity matrix, which corresponds to well-known brain regions. *** *p* < .001, ** *p* < .01, * *p* < .05.

Since the morphogen gradients have been established and their pattern was conserved during the development ^19^, we explored whether these genetic patterns correspond with those of the structural connectivity. We first identified the genes in AHBA that are strongly expressed in one end of the brain, according to a previous publication ^39^. We subdivided the connectopies equally into ten parts. Two sample t-tests were used to assess the difference in gene expression of these genes between the first and the last bin. We found expression of *FGF17* was significantly higher in the dorsal cluster than the ventral cluster (*t* = 27, *p* < .001), while the expression of *FOXG1* was significantly higher in the ventral (*t* = −25, *p* < .001). The same opposite patterns were observed between *FGF8* (rostral < caudal, *t* = −33, *p* < .001) and *PAX6* (rostral > caudal, *t* = 29, *p* < .001), and *SFRP1* (medial < lateral, *t* = −41, *p* < .001) and *WNT3* (medial > lateral, *t* = 99, *p* < .001) (Figure 5a). The number of subdivisions does not affect the final results (Figure S12).

**Figure 5.**
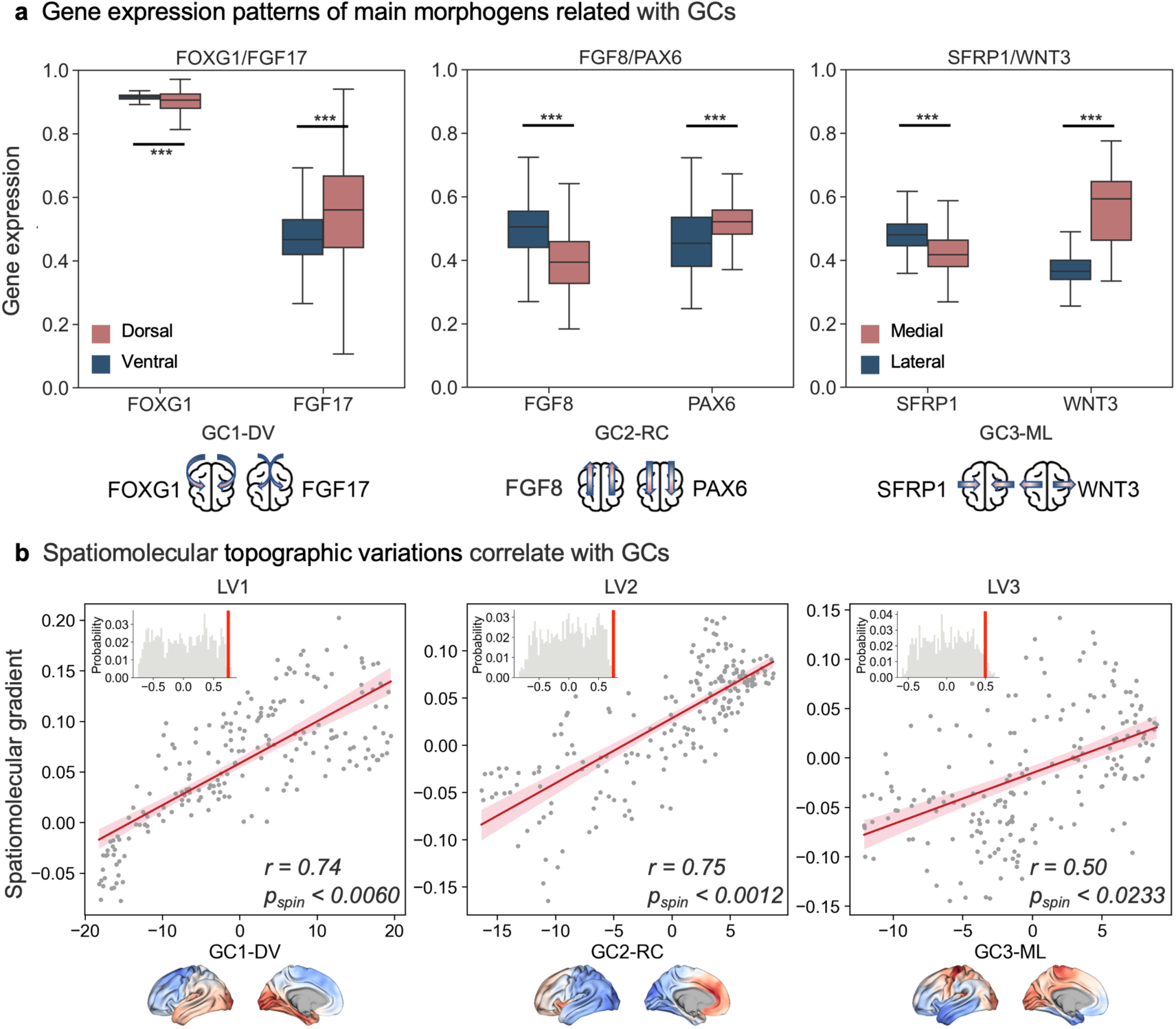
Consistency between global connectopies and morphogen gradients and spatiomolecular gradients. **(a)** The expression of morphogen genes that express the same pattern as any of the three global connectopies was significantly different between the two ends of the global connectopies (two-sample t-test, all *p* < .001). **(b)** The global connectopies significantly correlated with the spatiomolecular gradients, which maintained the same pattern as morphogenetic gradients during development (*r*_GC1-DV, LV1_ = 0.74, *p*_spin_ < .0060, FDR corrected; *r*_GC2-RC, LV2_ = 0.75, *p*_spin_ < .0012, FDR corrected; *r*_GC3-ML, LV3_ = 0.5, *p*_spin_ < .0233, FDR corrected). *** *p* < .001.

We also showed that the three global connectopies showed significant correlations with the spatiomolecular gradients ^19^ (*r*_GC1-DV, LV1_ = 0.74, *p*_spin_ < .0060, FDR corrected; *r*_GC2-RC, LV2_ = 0.75, *p*_spin_ < .0012, FDR corrected; *r*_GC3-ML, LV3_ = 0.5, *p*_spin_ < .0233, FDR corrected; Figure 5b). The spatiomolecular gradients were reported to remain in the same pattern as morphogenetic gradients during development. Note that the gradient values for the subcortex, cerebellum, and brainstem were excluded, and only data for the cortex were considered. Since the LV3 gradient showed the most variation and varied along a mediolateral and dorsoventral direction, as mentioned in ^19^, the *r*-value was lower compared with the former two but still significant, and the correlation between the first global connectopy was also moderate (*r*_GC1-DV, LV3_ = −0.36, *p*_spin_ < .05). These again suggested a potential genetic basis underlying the global connectopies.

Finally, we explored the roles of genes that contributed to global connectopies. Here we focused on the genes in the top 5% of each global connectopy (corrected for multiple comparisons). We found that a selection of genes contributed to more than one global connectopy, with 22 genes overlapping with all three (Figure 6a, right). All significantly associated genes overlapped with known morphogenetic genes identified previously ^39^ (hypergeometric test, *p* < .001; Figure 6b). Gene enrichment analyses using different gene sets showed that genes that contributed to only one global connectopy, especially GC2-RC, were involved with biological processes (Figure 6c). The top genes in each global connectopy, such as *FARSA*, *MPND*, *SYCP2*, *CUX1*, *ARHGDIG*, and *NUDT14*, are known to be associated with the regulation of transcription factor activity, morphogenesis, cell proliferation, and metabolic processes (Figure 6b) ^40–46^, while genes related to all three global connectopies, including *ADAMTSL1* and *TIAM1*, showed an obvious connection with various diseases (Figure 6b) ^47–49^. We found these genes significantly enriched in radial glial cells in prenatal samples (Figure 6c, left) and enriched in excitatory neurons after birth (Figure 6c, right). This was verified by the developmental pattern of these genes, which were expressed highly in the prenatal period in all lobes and that their expression decreased after birth (Figure 6d), suggesting that the patterns present in adulthood were nearly reached in the early stages of development. Digging deeper into specific biological processes, we found an enrichment of terms related to the regulation of transcription, metabolic process, morphogenesis, cellular development, and neuron projection (Figure 6e).

**Figure 6.**
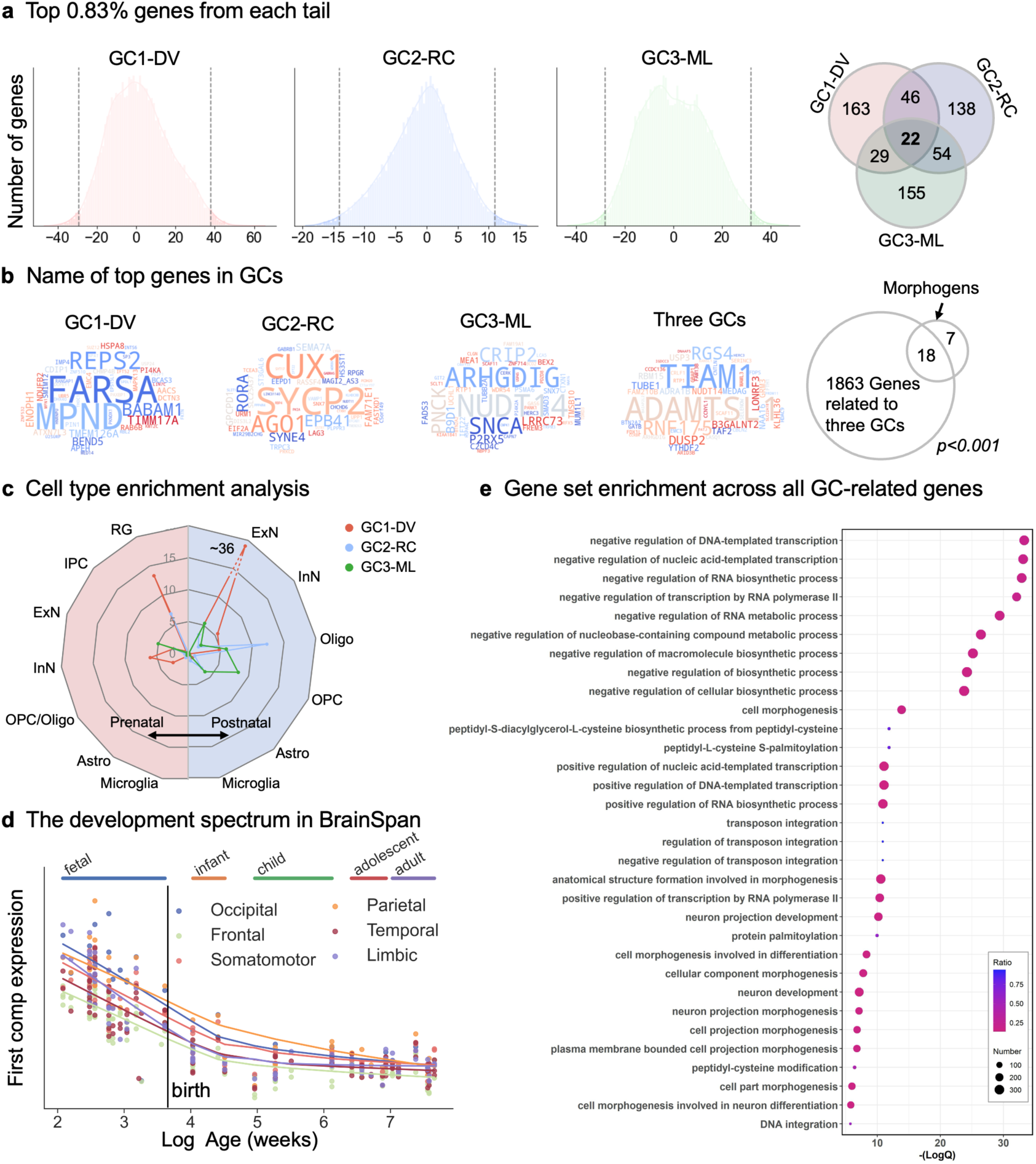
Gene enrichment analysis of connectopy-associated genes. **(a)** Distributions of connectopic weights across genes. Genes in the top 0.83% of either tail of the distribution (*n* = 130) were selected for further enrichment analysis. The Venn diagram showing the overlap of genes is shown on the right. **(b)** Top genes for unique connectopy and overlap of connectopies. **(c)** Cell type enrichment analysis was conducted on genes belonging to each GC using prenatal and postnatal datasets. These genes are significantly enriched in radial glial cells in prenatal samples and enriched in excitatory neurons after birth. **(d)** The development spectrum of gene expression for each brain macrostructure. Genes associated with GCs were expressed highly in the prenatal period and decreased after birth. **(e)** Gene set enrichment across all GC-related genes indicated terms related to the regulation of transcription, metabolic process, morphogenesis, cellular development, and neuron projection.

## Discussion

In the current study, we demonstrated that a low-dimensional, topological representation of brain connectivity exists and may share common space with gene expression despite the significant disparity in the numbers of genes and connections. We systematically analyzed the spatial topography of cerebral connectivity, i.e., the white matter tractogram, and identified three orthogonal global connectopies: the dorsoventral, rostrocaudal, and mediolateral connectopy across the cerebral cortex, which is represented in the spatial arrangement of long-range white matter tracts. Moreover, the global connectopies were observed to be aligned with the gradients of morphogens identified during embryonic brain development and with genetic topography and spatiomolecular gradients. Our findings demonstrate the crucial role that comprehending the connectivity topographies and their genetic constraint plays in understanding the underlying principles that shape the human brain organization.

### Global connectopies characterize the arealization consistent with genetic gradients

Our findings showed that the three global connectopies derived from the white matter partly characterized the embedding of connectivity architecture underlying the cortex in a low dimensional space organized along the dorsoventral, rostrocaudal, and mediolateral axes, which closely resemble the principal axes of brain development ^19,50^ and the chemotactic gradients of early embryogenesis ^51,52^. From a developmental perspective, both the encoding of chemical gradients by morphogens and the spatial expression patterns of all genes during the formation of the mature brain, along with recent research on phenotypic mapping facilitated by twin studies ^1–3^, indicate that gradient-based organizational principles may serve as one of the fundamental rules in the precise formation of gene-encoded connections and the establishment of functional cortical regions. Meanwhile, genes provide the initial blueprint that guides differentiation and the directional migration of neurons through the subsequent formation of different types of gradients of transcription factors, morphogens, etc., and, in parallel, the precise trajectory and projection of axons ^53^. However, the reality is likely somewhere in between that a few dozen molecules (rather than a few or a thousand) enact pattern formation rules to increase the probability that neurons make the correct connections. The global connectome organization might partly be governed by simple genetic rules. The scaffolding rule guides the exact formation of neural circuits ^54^, wherein genes might not drive the precise details of complex circuit diagrams but have the potential to influence their organizational rules during development ^22^. Here, the observed global connectopies could be one of the possible rules, consistent with the argument that anatomical connections could characterize the arealization and show consistency with cortical patterns driven by genetic profiles ^10,55,56^.

The dorsoventral and rostrocaudal genetic topographies (Figure 4) observed in our study suggest genetic influences on cortical morphology where regional differentiation is controlled by genes expressing a graded pattern ^1,2,57^. More interestingly, we not only observed a high correspondence between gene topology and our global connectopies but also found that modules identified using global connectopies closely resemble that obtained using genetic correlation ^2^. This correspondence indicates the presence of the same graded pattern across the cortex at the genetic and connectional levels, and they are consistent with the gradation of architectonic features. The connectivity between cortical layers is related to their architectonic features and may provide evidence for the role of connections in the initial scaffolding for cortical organization. Previous research found that cortical patterning creates segregated areas with different functions via cell differentiation and migration. These segregated areas specifically occur along molecular gradients representing the patterned expression of morphogens and transcription factors ^17,58^, which were previously established and found to radiate along the rostrocaudal, dorsoventral, and mediolateral axes of the neural compartment ^59^. These are conserved to maturity ^19^ and are consistent with our global connectopies (Figure 5b). At maturity, the morphogen gradients are replaced by a more distinct and discrete regional gene expression ^39^. Under the effect of morphogens secreted in the patterning centers with absolute positional information, numerous transcription factors are expressed in a graded manner. For example, the anterior expression of *FGF8* suppresses the posteriorly expressed transcription factors, resulting in their lower anterior but higher posterior expression ^60^. *FGF8* was also reported to stimulate the proliferation of cortical progenitor cells, thus regulating neurogenesis and arealization by thalamic axons ^61^. Similarly, manipulation of the transcription factor *EMX2* by genetic knockout resulted in finding higher expression posteromedially, as well as finding that a lower expression anterolaterally causes an expansion of the frontal and lateral regions at the expense of the visual cortex in mice ^62^. These transcription factors are believed to determine the areal fate and the expression of cell-surface molecules, thus determining the topographic organization of synaptic inputs and outputs ^1,20^ and directing axonal outgrowth ^21^. They significantly overlapped with the genes related to the identified global connectopies, which exhibited distinct developmental patterns and were especially involved in biological processes, including regulation of transcription, metabolic process, morphogenesis, cellular development, and neuron projection (Figure 6). In addition, the genes associated with all the global connectopies were found to be more related to brain diseases. This finding is consistent with the observation that disruptions in forming early developmental gradients through mutations to gradient-associated genes can cause severe developmental disorders ^19,63^. These associations demonstrated that global connectopies and their underlying genetic basis were essential to the development and function of the brain.

Many existing studies of thalamocortical and corticothalamic projections have identified specific receptors and ligands, such as the Eph receptor tyrosine kinases and their ephrin ligands, that are required to establish proper patterns of afferent and efferent cortical connectivity with subcortical structures ^64,65^. However, whether these molecules play a similar role in instructing intracortical connectivity within the developing cortex is unclear. Relevant studies in this regard, such as *in-utero* electroporation studies ^66^ and a study during primate development ^67^, suggest that directed axonal growth and target selection play a role in forming association projections between specific cortical areas. Moreover, according to the radial unit hypothesis of cortical development, radial glial cells in the embryonic brain facilitate the generation, placement, and allocation of neurons in the cortex and regulate how they wire up ^68,69^. The radial glial scaffold might act as a “highway” for axons, facilitating their growth and ensuring their proper trajectories. Nevertheless, the factors that drive the competence of specific axons to grow into selected gray matter regions are unknown, but molecular interactions between axons and recipient cortical areas may be involved ^53^. Here, our current findings only identified associations between gene gradients and global connectopies in adult brains. Although we also did an integrated lifespan analysis of genes that appear to contribute to the connectopies, providing possible correlations, further studies are needed to explain the dynamics and causality during early development directly. In future studies, the progress of new technologies ^70,71^ should be particularly helpful here and hopefully enable a clearer explanation of these relationships.

### Methodological consideration and future work

Several technical and methodological limitations must be acknowledged in the current work. The first and most direct concern is the accuracy of mapping structural connections using diffusion tensor imaging. However, diffusion tractography has been used as an irreplaceable tool to identify white matter across the brain *in vivo* and non-invasively, with certain limitations imposed by the tractography, such as gyral bias, which can cause false positive results ^72^. In the future, joint MRI and microscopy data analysis may help solve limitations ^73–75^. Alternatively, cutting-edge invasive neuro-techniques used in other mammalian cortexes, such as Cre-drivers for cortical projection mapping ^76^ or polarised light imaging ^77^, could be used at the mesoscale level of resolution to study whether such organizational axes are conserved across scales and species. The thresholding of the tractogram covariance and ignorance of negative edges should be considered in the context of diffusion map embedding, which may convey potential interacting anatomical connectivity profiles but lack accommodation in the current method. Distance effect also needs to be considered when mapping the structural connectivity pattern, but the consistent genetic topography of the three global connectopies suggests that they are not distance-dependent. Interestingly, a recent study challenged traditional views and showed that the geometry of the brain had a more fundamental role in shaping its function than interregional connectivity ^78^. Although the association between cortical geometry and brain function should be recognized, the interacting role of connectopies and geometry deserves further attention to reveal the function realization of the developing human brain, especially the neonate brain. The underlying genetic drivers should also be considered to explore the causal relationship between them. Secondly, the parcellation guided by the three global connectopies is very coarse compared with several fine-grained brain atlases that have been delineated using the gradient approach ^79,80^. We did not utilize these more fine-grained parcels because we believe they are more likely to be shaped by local gradients compared with the three global connectopies, which characterize the global pattern across the brain.

In this study, we mainly focused on the structural connectivity patterns shown on the cortex and brought functional emergence into the discussion. More thorough microscale biological foundations and physiological processes were not explored here, but in future studies, they may aid in interpreting our findings about macroscale structural patterns. Since the spatiomolecular gradients expressed in childhood and adulthood have been established and shown to be consistent ^19^, an immediate goal will be to explore the transition in structural patterns throughout development. Whether these global connectopies are conserved across species could be another perspective to study when looking into the organization of the brain. We also need animal studies to validate or study the causal relationships between the genes, connectivity, and parcellations, which will reduce the impact of mismatch of ages and other factors resulting from relating HCP young adult diffusion MRI connectopies to AHBA-based gene expression information. Although the AHBA is the densest sample transcriptomic dataset of the human brain to date, it comes with the above-mentioned limitations, as well as the gap between post-mortem changes in gene expression level and *in vivo* features, making our results a matter of caution. With the development of RNA sequencing technology, which enables rapid profiling and deep investigation of the transcriptome, more genes could be detected and provide valuable information ^81^. In addition, the observed organizing principles in the current work may also be used to guide the engineering of three-dimensional cortical spheroids and to comprehend the human-specific aspects of the neural circuit assembly ^82,83^.

### Conclusions

To conclude, we explored how genes link the accurate formation of connections despite significant differences in their numbers. Our research uncovered three determinant spatial trends across the cerebral cortex, aligning with the dorsoventral, rostrocaudal, and mediolateral axes, which we identified as global connectopies. These global connectopies were found to be aligned with the gradients of morphogens identified during embryonic brain development and with genetic topography and spatiomolecular gradients. Our findings revealed the existence of a low-dimensional, topological representation of brain connectivity that may share a common space with gene expression, providing a consistent characterization of the cortical arealization. We hope our discoveries will offer valuable insights into studying brain structure and function.

## Materials and Methods

### Data and preprocessing

#### Data collection

We used a publicly available dataset containing 100 unrelated subjects (HCP-U100) (46 males; mean age, 29.11±3.67; age range, 22-36) provided by the Human Connectome Project (HCP) database ^32^ (http://www.humanconnectome.org/). All the scans and data from the individuals included in the study had passed the HCP quality control and assurance standards.

The scanning procedures and acquisition parameters were detailed in previous publications ^84^. In brief, T1w images were acquired with a 3D MPRAGE sequence on a Siemens 3T Skyra scanner equipped with a 32-channel head coil with the following parameters: TR = 2400 ms, TE = 2.14 ms, flip angle = 8°, FOV = 224×320 mm^2^, voxel size = 0.7 mm isotropic. Diffusion data were acquired using single-shot 2D spin-echo multiband echo planar imaging on a Siemens 3 Tesla Skyra system (TR = 5520 ms, TE = 89.5 ms, flip angle = 78°, FOV = 210×180 mm). These consisted of three shells (b-values = 1000, 2000, and 3000 s/mm^2^), with 90 diffusion directions isotropically distributed among each shell and six b = 0 acquisitions within each shell, with a spatial resolution of 1.25 mm isotropic voxels.

#### Image preprocessing

The human T1w structural data had been preprocessed following the HCP’s minimal preprocessing pipeline ^84^. In brief, the processing pipeline included imaging alignment to standard volume space using FSL, automatic anatomical surface reconstruction using FreeSurfer, and registration to a group average surface template space using the multi-modal surface matching (MSM) algorithm ^85^. Human volume data were registered to the Montreal Neurological Institute (MNI) standard space, and surface data were transformed into surface template space (fs_LR).

The diffusion Images were processed using FDT (FMRIB’s Diffusion Toolbox) of FSL. The main steps are normalization of the b0 image intensity across runs and correction for echo-planar imaging (EPI) susceptibility, eddy-current-induced distortions, gradient-nonlinearities, and subject motion. DTIFIT was then used to fit a diffusion tensor model. The probability distributions of the fiber orientation distribution were estimated using Bedpostx.

Next, skull-stripped T1-weighted images for each subject were co-registered to the subject’s b0 images using FSL’s FLIRT algorithm. Then nonlinear transformations between the T1 image and the MNI structural template were obtained using FSL’s FNIRT. Based on concatenating these, we derived bi-directional transformations between the diffusion and MNI spaces.

### Connectopy mapping

#### Tractography and connectivity blueprints

To map the whole-brain connectivity pattern, we performed probabilistic tractography using FSL’s probtrackx2 accelerated by using GPUs ^86,87^. Specifically, the white surface was set as a seed region tracking to the rest of the brain with the ventricles removed and down-sampled to 3 mm resolution. The pial surface was used as a stop mask to prevent streamlines from crossing sulci. Each vertex was sampled 5000 times (5000 trackings) based on the orientation probability model for each voxel, with a curvature threshold of 0.2, a step length of 0.5 mm, and a number of steps of 3200. This resulted in a (*whole-surface vertices*) × (*whole-brain voxels*) matrix for further analysis. We checked the preprocessing images and tractography results of each subject to ensure the accuracy of our analysis. Specifically, the reconstructed surfaces were visually inspected, and transforms between different spaces were checked carefully. Tractography results were also visually checked to see if any streamline crosses the sulci.

We additionally created connectivity blueprints following previous work ^36^, which had been used to characterize connectivity patterns of brain regions or cortical vertices. Specifically, we first reconstructed 72 fiber bundles using a pre-trained deep-learning model, TractSeg ^35^, and down-sampled the resulting tract masks to 3 mm resolution, yielding a (*tracts*) × (*whole-brain voxels*) matrix. Then, the connectivity blueprints were generated by the product of this tract matrix, and the whole-brain connectivity matrix was obtained and normalized as described in the Tractography section. The columns of the resulting (*tracts*) × (*whole-surface vertices*) matrix showed the connectivity distribution pattern of each cortical vertex, while the rows revealed the cortical termination patterns of the tracts.

#### Tractogram covariance analysis

We calculated the tractogram covariance (TC) matrix to characterize the structural connectivity similarity profile. The vertex profiles underwent pairwise Pearson correlations, controlling for the average whole-cortex profile. For a given pair of vertices, *i* and *j,* TC was calculated as

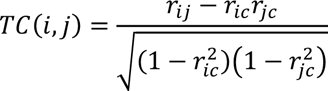

in which *r_ij_* was the Pearson correlation coefficient of the structural connectivity profile at vertices *i* and *j*, *r_ic_* the correlation of the structural connectivity profile at vertex *i* with the average connectivity profile across the whole cortex, and *r_jc_* the correlation of the structural connectivity profile at vertex *j* with the average connectivity profile across the entire cortex. A symmetric ∼30k×30k TC matrix was produced for each subject. Then, the TC matrices of all subjects were averaged separately for the left and right hemispheres to get a group-level TC matrix for each hemisphere. In line with a previous study ^88^, the TC matrix was proportionally thresholded at 90% per row, with elements above zero retained to remove negative connections.

#### Connectopy decomposition

The TC matrix was transformed into a non-negative square symmetric affinity matrix using a cosine affinity kernel. Then, diffusion map embedding was implemented to identify the principal gradient components ^33^. Diffusion map embedding is a nonlinear manifold learning technique that maps cortical gradients ^27^. Along these gradients, cortical vertices with similar connectivity profiles are embedded along the axis. Diffusion map embedding is relatively robust to noise and less computationally expensive than other nonlinear manifold learning techniques.

Global connectopies were computed separately for the left and right hemispheres with two key parameters: *α* controls the influence of the density of sampling points on the manifold, and *t* controls the scale of the eigenvalues. Here, we set *α*=0.5 and *t*=0 as recommended ^27^. The amount of explained variance was assessed, and the first three global connectopies were chosen to map onto the cortical surface for further analysis.

Individual global connectopies were calculated for each subject and aligned to the group GC with Procrustes rotation, which matches the order and direction of the global connectopies without scaling. The application of Procrustes rotation allows for comparison between individuals.

#### Projection of the global connectopies onto the white matter voxels

We used general linear models (GLM) to project the global connectopies onto the white matter ^34^, exploring the contribution of each white matter voxel to the global connectopies. Specifically, we used the global connectopies as a dependent variable and the structural connection matrix as an independent variable. The resultant weights indicate the contribution of each white matter voxel. Note that the structural connection of each subject was obtained from probabilistic tractography in individual diffusion space. We registered each to MNI standard space and averaged, resulting in a group-level structural connection matrix.

#### Projection of the connectopies onto the white matter tracts

To investigate how the global connectopies are related to the underlying white matter tracts, we made a tract projection map using the tract mask generated by TractSeg. Seventy-two tracts identified by TractSeg were grouped into five types for analysis: association tracts, commissural tracts, projection tracts, thalamo tracts, and striato tracts (Table S1). Using the connectivity blueprints we built before, we extracted the projection on the surface of each tract and calculated the mean value and variance using connectopy values of tract projection on the surface, thus revealing the relationship between the tracts and global connectopies. Due to the absence of projection on the surface, we excluded the superior cerebellar peduncle (SCP), middle cerebellar peduncle (MCP), and inferior cerebellar peduncle (ICP) from the study. We additionally removed Corpus Callosum – all because several segments were also reconstructed by TractSeg, i.e., CC_1 to CC_7.

#### Visualization of connectopies

To inspect the global connectopies together intuitively, we plotted every two global connectopies in a 2D space and assigned them colors based on their positions. Then we generated RGB values for each vertex by normalizing three global connectopy values and mapped them on the surface, showing the dominant global connectopy at the vertex. Specifically, for the first gradient, values of all vertices were normalized from zero to one, resulting in a value of the red channel. The other two were conducted the same.

### Hierarchical organization analysis

We implemented hierarchical clustering using global connectopies to test if the global connectopies could provide a descriptor of arealization. At the first level, the vertices were sorted in descending order from the first global connectopy. Brain regions were partitioned into two subregions based on the sign of the connectopy value. Furthermore, based on the positivity and negativity of the second and third global connectopies, the whole cortex was partitioned into eight modules.

### Gene analysis

#### Genetic correlation analysis of thickness and surface area

We selected 194 twins from the HCP database (102 monozygotic and 92 dizygotic pairs; mean age, 29.96±2.96; age range, 22-36). The cortical surface was reconstructed to measure the cortical thickness and surface areas at each surface location using FreeSurfer in the fsaverage3 template to reduce computation time.

In a classical twin study of sets of MZ and DZ twins, four latent factors could account for the variance of any phenotype: additive genetic effects (A); non-additive genetic effects, including dominance (D); common or shared environmental effects (C); and non-shared or individual-specific environmental effects. Because of the minimal difference between the A and D effects, we fitted bivariate ACE models to compute the genetic correlations of the cortical thickness and surface area between two vertices of the cortex. Thus, the genetic covariance was calculated decomposed from the total phenotypic correlation. The analyses were performed using OpenMx ^89^.

Then we implemented diffusion map embedding on the genetic correlation matrix and compared it with the global connectopies. Genetic patterning results were obtained from previous studies ^1,2^, which parcellated the cortex into four and twelve clusters. The parcellation results were compared with the modules derived from hierarchical clustering using the global connectopies.

#### AHBA data and preprocessing

Regional microarray gene expression data were obtained from 6 postmortem brains (1 female, ages 24-57) provided by the Allen Human Brain Atlas (AHBA, https://human.brain-map.org/). The data were processed using the abagen toolbox (version 0.1.1; https://github.com/rmarkello/abagen) ^90^. First, the microarray probes were reannotated ^91^, and the probes that did not match a valid Entrez ID were excluded. Then, the probes were filtered based on their expression intensity relative to background noise, and the probes with the maximum summed adjacency for representing the corresponding gene expression were kept, yielding 15633 genes that corresponded to more than one probe. Last, we resampled the output gene expression map in fs5 space to fsaverage_LR32k space for subsequent study.

#### Analysis of morphogen gradient-related genes

We further selected morphogen genes expressing the same pattern as any of the three global connectopies in AHBA ^39^. Each global connectopy was subdivided equally into ten parts, and two sample t-tests was used to assess the difference in gene expression of selected genes between the first and the last bin ^92^.

#### Correspondence to spatiomolecular gradients

Researchers recently reported three spatiomolecular gradients that retained the same pattern as morphogenetic gradients during their development. These gradients varied along three spatially embedded axes ^19^. We only used data from the left hemisphere because few samples in the AHBA were obtained from the right hemisphere. We replicated these molecular gradients at the ROI level with the “Schaefer-400” parcellation scheme ^80^ and explored the correlation with the three global connectopies in this study.

#### Gene enrichment analysis

We used a prediction framework, partial least squares regression (PLSR), to determine the covariance between gene expression and global connectopies. Then we conducted 1000 bootstrapping to estimate the error of the weight of each gene. The normalized weight of each gene was denoted as the weight divided by the estimated error ^93^.

Genes were selected as the top and bottom 0.83% genes (*n* = 260) of each global connectopy’s gene list, corrected for both tails and all three global connectopies for multiple comparisons. Each gene set combining both tails underwent cell-type enrichment analysis on the filtered genes using CellGO (http://www.cellgo.world) ^94^. Single-cell datasets collected from prenatal ^95^ and postnatal sample were used ^96^. Seven major classes of cells were provided in prenatal sample, including radial glial cells (RG), intermediate progenitor cells (IPC), excitatory neurons (ExN), inhibitory neurons (InN), oligodendrocyte precursor cells/oligodendrocytes (OPC/Oligo), astrocytes (Astro), and microglia cells (Microglia). Six major classes of cells were provided in postnatal sample, including ExN, InN, Oligo, OPC, Astro, and Microglia. The enrichment P-values of the cell types resulting from the submitted genes were based on the Kolmogorov-Smirnov (K-S) test in CellGO.

The filtered genes extended across the three global connectopies were also submitted to the gene ontology enrichment analysis. ToppGene (https://toppgene.cchmc.org/), which contains the whole list of AHBA genes as the background gene set, was used to conduct the analysis. The following term categories were assessed: GO: Molecular Function, GO: Biological Process, GO: Cellular Component, Pathway, and Disease.

We collected gene expression data for all time points in the BrainSpan dataset to determine the developmental pattern of gene expression ^97^. Only genes associated with all three global connectopies were selected. Non-negative matrix factorization was used to derive the gene expression components for each macrostructure in the brain.

### Robustness analyses

Complementing our main analysis on calculating global connectopies, we examined the effect that may caused by age, sex, and brain size. For age, we divided subjects into three groups (age 20-25: *n* = 17; age 25-30: *n* = 40; age 30-35: *n* = 43). For sex, we divided the subjects into males (*n* = 46) and females (*n* = 54). For brain size, we extracted each HCP subject’s total intracranial volume (TIV) from the Freesurfer output. Moreover, we divided into five groups (TIV 1.0-1.2: *n* = 3; TIV 1.2-1.4: *n* = 12; TIV 1.4-1.6: *n* = 38; TIV 1.6-1.8: *n* = 40; TIV 1.8-2.0: *n* = 7; [*10^6^ mm3]). We calculated the correlation of the tractogram covariance matrix between groups.

We further derived the geodesic distance matrices on the mid-thickness surface of the human brain, calculated the gradients, and computed the correlation with global connectopies to test the effect by geodesic distance. We also remove short-range connections in the tractogram covariance matrices. We removed the value if the geodesic distance of the two vertices was less than 10mm, 20mm, and 30mm, respectively, and calculated the global connectopies again.

## Data availability

Data from the Human Connectome Project can be downloaded at https://db.humanconnectome.org/. The human gene expression data are available in the Allen Brain Atlas (https://human.brain-map.org/static/download). Data from BrainSpan can be downloaded at https://www.brainspan.org/.

## Code availability

The HCP pipeline can be found at https://github.com/Washington-University/HCPpipelines. The neuroimaging preprocessing software used for the other datasets is freely available (FreeSurfer v6.0, http://surfer.nmr.mgh.harvard.edu/, and FSL v6.0.5, https://fsl.fmrib.ox.ac.uk/fsl/fslwiki). The gene processing pipeline is available (abagen, https://github.com/rmarkello/abagen), and gene enrichment analysis is conducted at https://toppgene.cchmc.org/. The brain maps were presented using BrainSpace (https://brainspace.readthedocs.io/) and Connectome Workbench v1.5.0 (https://www.humanconnectome.org/software/connectome-workbench). The tracts were visualized using BrainNet Viewer v1.7 (https://www.nitrc.org/projects/bnv/). Python code to reproduce the analyses and figures is available at https://github.com/FANLabCASIA/GC.git (will make this public upon acceptance).

## Acknowledgments

This work was partially supported by STI2030-Major Projects (Grant No. 2021ZD0200200), the Natural Science Foundation of China (Grant Nos. 82072099, 82202253, 62250058), and the China Postdoctoral Science Foundation (2022M722915). Data were provided in part by the Human Connectome Project, WU-Minn Consortium (Principal Investigators: David Van Essen and Kamil Ugurbil; 1U54MH091657) funded by the 16 NIH Institutes and Centers that support the NIH Blueprint for Neuroscience Research; and by the McDonnell Center for Systems Neuroscience at Washington University. The authors appreciate the English language and editing assistance of Rhoda E. and Edmund F. Perozzi, PhDs.

## Author Contributions

DL, CC, LF designed the research; DL performed the experiments; YW, LM, SW, YL, HW, YaW, CTE, CG, LC, LF, CC contributed new analytic tools; DL analyzed the data; DL, YW, LF, CC wrote the paper; and LF, CC, SBE, CHC, ZY, YZ contributed analytical expertise, theoretical guidance, paper revisions, and informed interpretation of the results.

