## Supplemental Table S1, Supplemental Figure S1-S12 for "Topographic Axes of Wiring Space Converge to Genetic Topography in Shaping Human Cortical Layout"

### Supplementary Materials

**This file includes**

Table S1

Figures S1 to S12

#### Supplementary tables

| Category | Fiber | Abbreviation | Left/Right? |
|---|---|---|---|
| Association | Arcuate fascicle | AF | Yes |
| | Cingulum | CG | Yes |
| | Middle longitudinal fascicle | MLF | Yes |
| | Inferior occipito-frontal fascicle | IFO | Yes |
| | Inferior longitudinal fascicle | ILF | Yes |
| | Optic radiation | OR | Yes |
| | Parieto-occipital pontine | POPT | Yes |
| | Superior longitudinal fascicle I | SLF_I | Yes |
| | Superior longitudinal fascicle II | SLF_II | Yes |
| | Superior longitudinal fascicle III | SLF_III | Yes |
| | Uncinate fascicle | UF | Yes |
| Projection | Corticospinal tract | CST | Yes |
| | Fronto-pontine tract | FPT | Yes |
| | Inferior cerebellar peduncle | ICP | Yes |
| | Middle cerebellar peduncle | MCP | No |
| | Superior cerebellar peduncle | SCP | Yes |
| Commissural | Fornix | FX | Yes |
| | Commissure anterior | CA | No |
| | CC rostrum | CC_1 | No |
| | CC genu | CC_2 | No |
| | CC rostral body | CC_3 | No |
| | CC anterior midbody | CC_4 | No |
| | CC posterior midbody | CC_5 | No |
| | CC isthmus | CC_6 | No |
| | CC splenium | CC_7 | No |
| | Corpus Callosum – all | CC | No |
| Thalamo | Anterior thalamic radiation | ATR | Yes |

| | | | |
|---|---|---|---|
| | Superior thalamic radiation | STR | Yes |
| | Thalamo-prefrontal | T_PREF | Yes |
| | Thalamo-premotor | T_PREM | Yes |
| | Thalamo-precentral | T_PREC | Yes |
| | Thalamo-postcentral | T_POSTC | Yes |
| | Thalamo-parietal | T_PAR | Yes |
| | Thalamo-occipital | T_OCC | Yes |
| Striato | Striato-fronto-orbital | ST_FO | Yes |
| | Striato-prefrontal | ST_PREF | Yes |
| | Striato-premotor | ST_PREM | Yes |
| | Striato-precentral | ST_PREC | Yes |
| | Striato-postcentral | ST_POSTC | Yes |
| | Striato-parietal | ST_PAR | Yes |
| | Striato-occipital | ST_OCC | Yes |

**Tables S1. 72 fibers reconstructed by TractSeg.**

### Supplementary figures

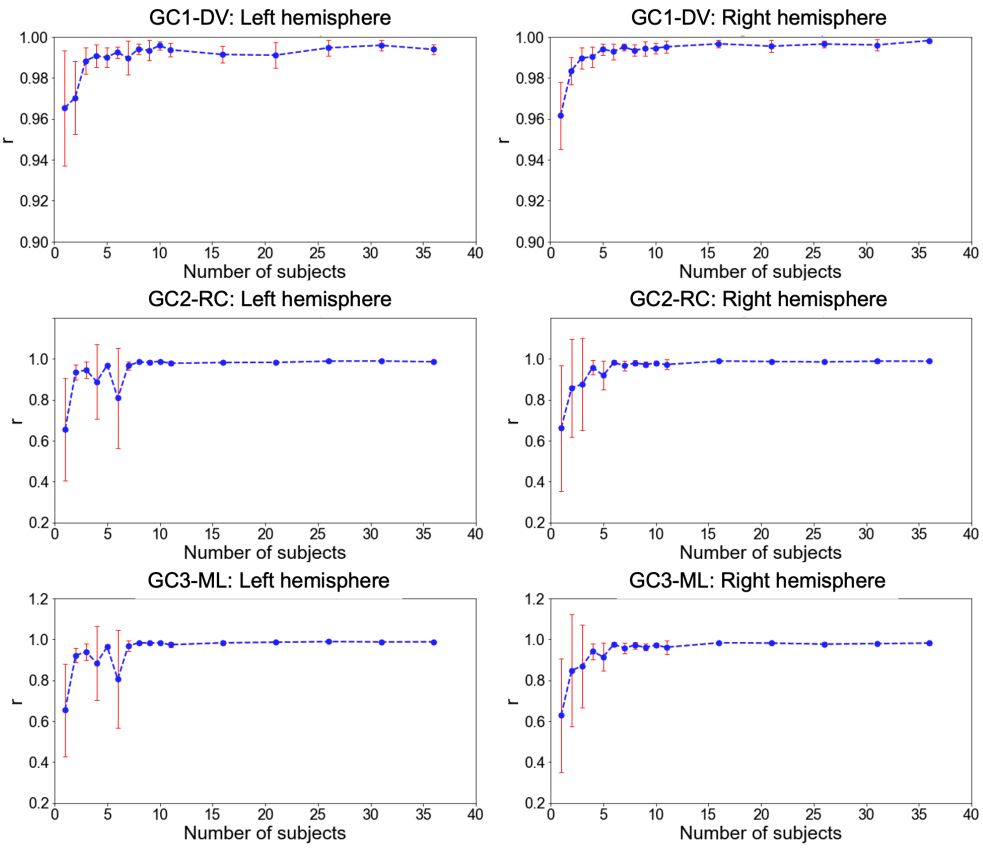


**Figure S1. Stability of the three global connectopies patterns across the subjects.** The correlation coefficient was greater than 0.9 when the number of subjects reached ~10.

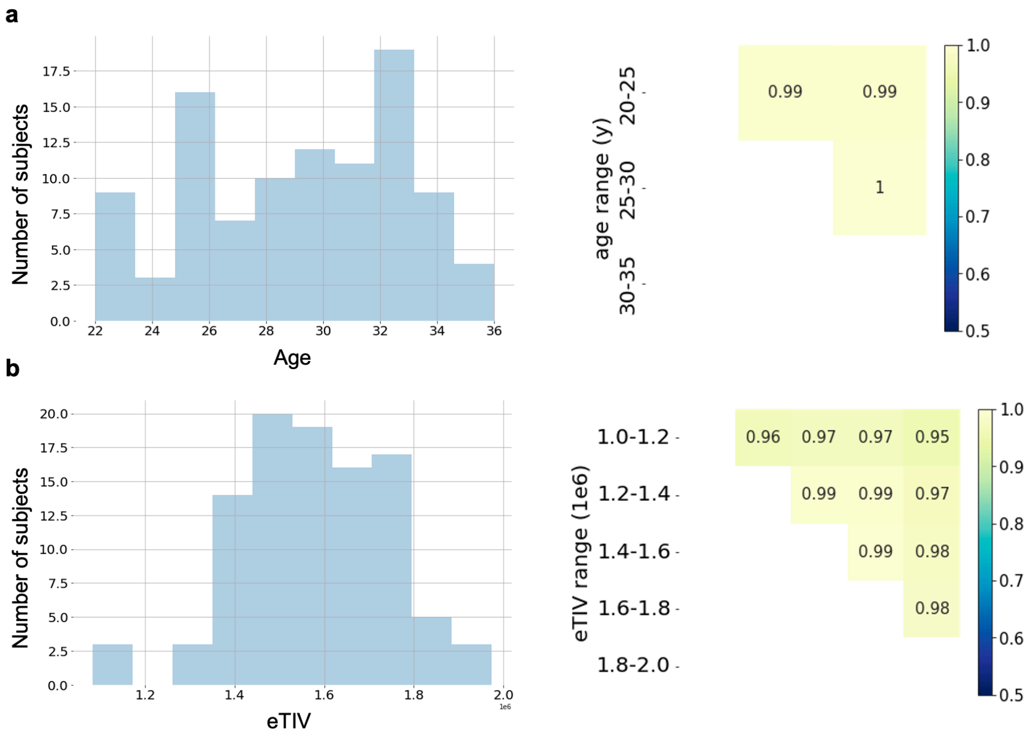


**Figure S2. The relationship between tractogram covariance and age, sex, and brain size. (a)** We divided subjects into three groups (age 20-25: $n$ = 17; age 25-30: $n$ = 40; age 30-35: $n$ = 43) and found that the tractogram covariance matrix among the three groups showed very high similarity. **(b)** We extracted each HCP subject's total intracranial volume (TIV) from the Freesurfer output. Furthermore, we divided into five groups (TIV 1.0-1.2: $n$ = 3; TIV 1.2-1.4: $n$ = 12; TIV 1.4-1.6: $n$ = 38; TIV 1.6-1.8: $n$ = 40; TIV 1.8-2.0: $n$ = 7; [*$10^6$ $mm^3$]) and calculated the tractogram covariance matrix. We found that the three groups showed very high similarity. We also calculated the correlation between the male group ($n$ = 46) and the female group ($n$ = 54) and found the correlation very high ($r$ = 0.99).

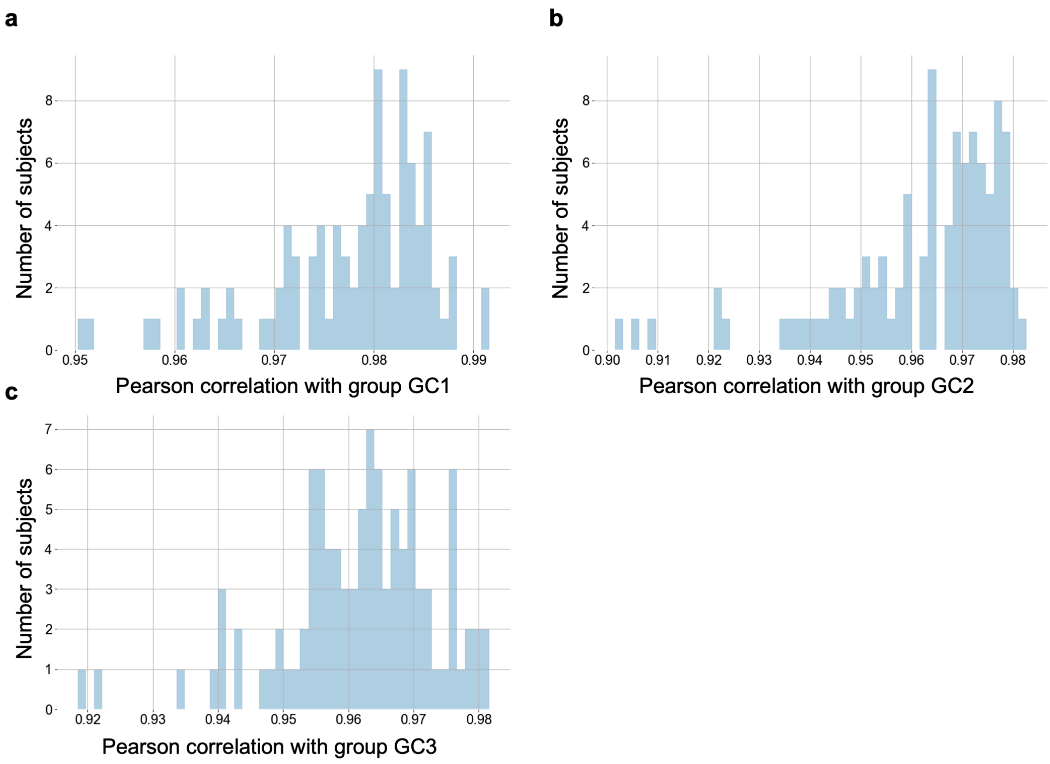


**Figure S3. Individual global connectopies were aligned after Procrustes alignment.** We applied Procrustes alignment on the individual's three global connectopies, i.e., **(a)** GC1-DV, **(b)** GC2-RC, and **(c)** GC3-ML. The similarity among individuals is very high (all $r > 0.9$).

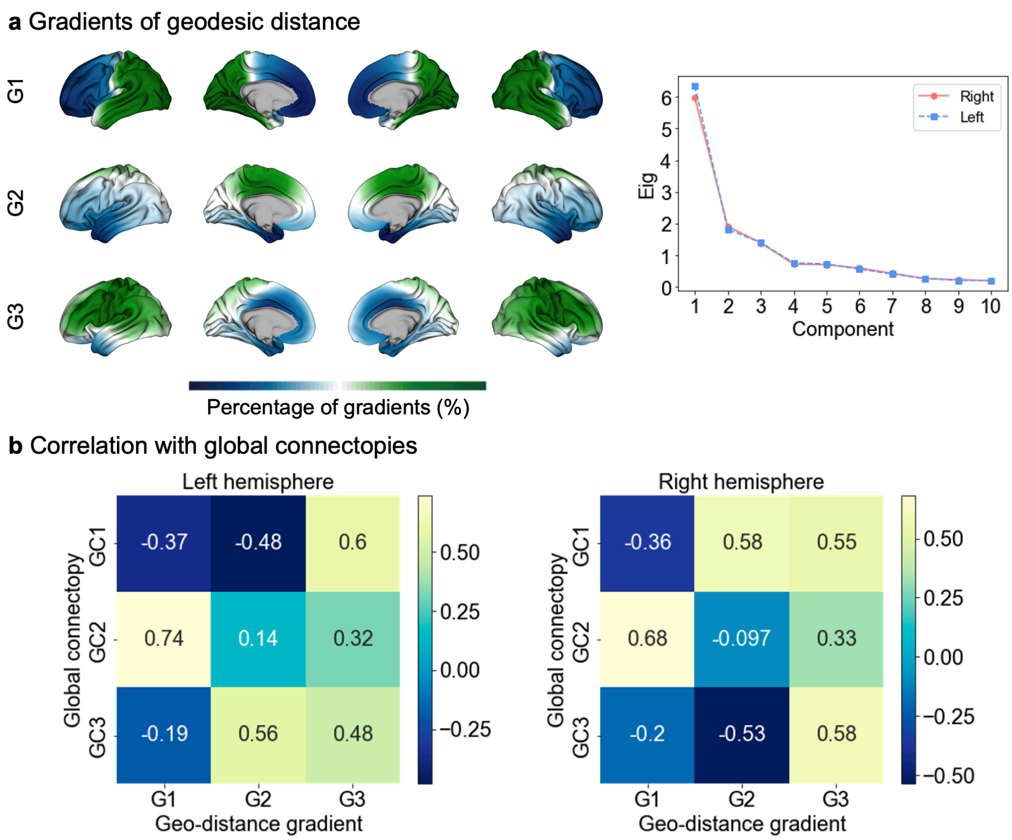


**Figure S4. Gradients are calculated by the geodesic distance matrix and its correlation with global connectopies. (a)** We derived the geodesic distance matrices on the mid-thickness surface of the human brain and calculated the gradients, which showed dissimilar patterns with our global connectopies. A scree plot describing variance is shown on the right. **(b)** Correlation between geodesic distance gradient and global connectopies.

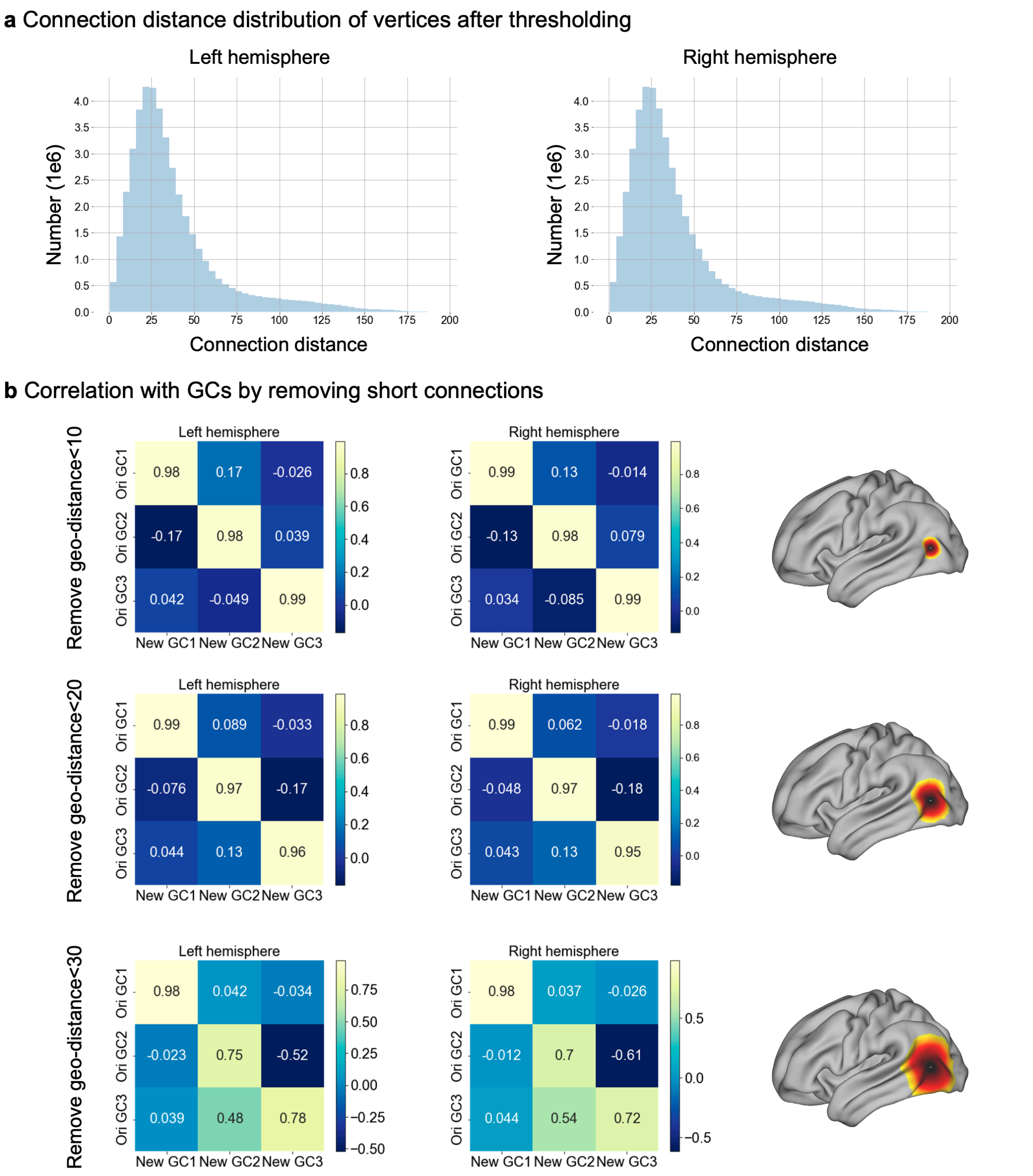


**Figure S5. Global connectopies are beyond geodesic distance and short-range connections. (a)** Connection distance distribution of the vertex after the thresholding step. Many long-distance connections remained after thresholding. **(b)** Global connectopies are calculated by removing the short-range connection and its correlation with global connectopies. We removed the value if the geodesic distance of the two vertices was less than 10mm, 20mm, and 30mm, respectively, and calculated the global connectopies again. We found that they showed very similar results to our original main results.

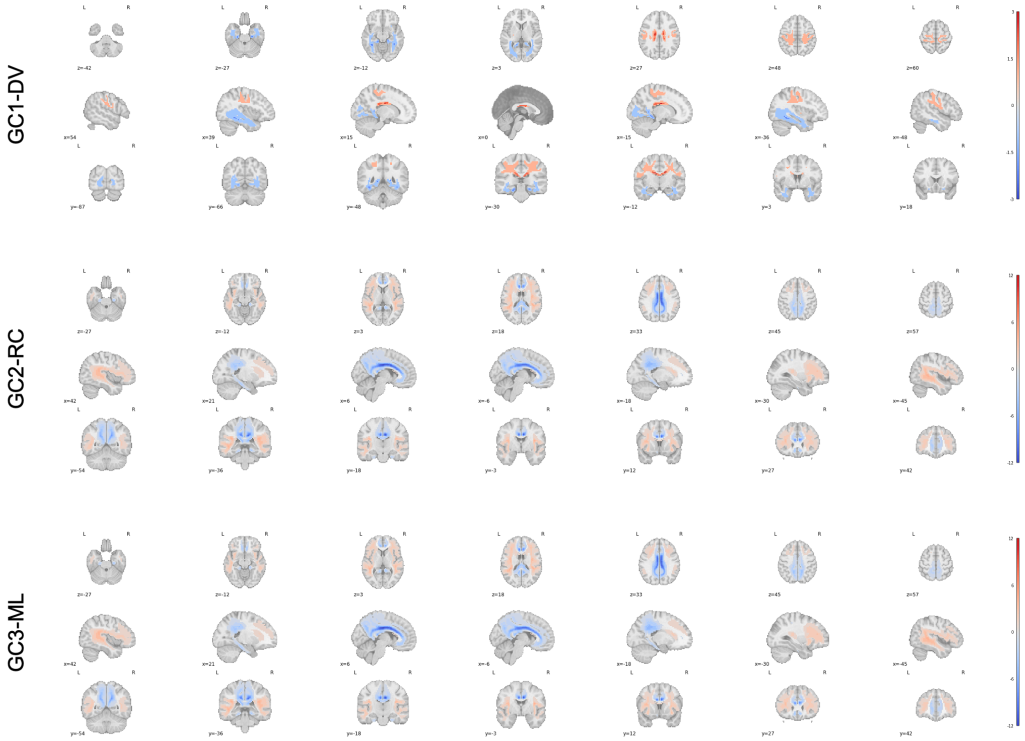


**Figure S6. Voxel's contribution to the global connectopies was calculated using the GLM method.** We used general linear models (GLM) to project the global connectopies onto the white matter, exploring the contribution of each white matter voxel to the global connectopies. Specifically, we used the global connectopies as a dependent variable and the structural connection matrix as an independent variable. The resultant weights indicate the contribution of each white matter voxel.

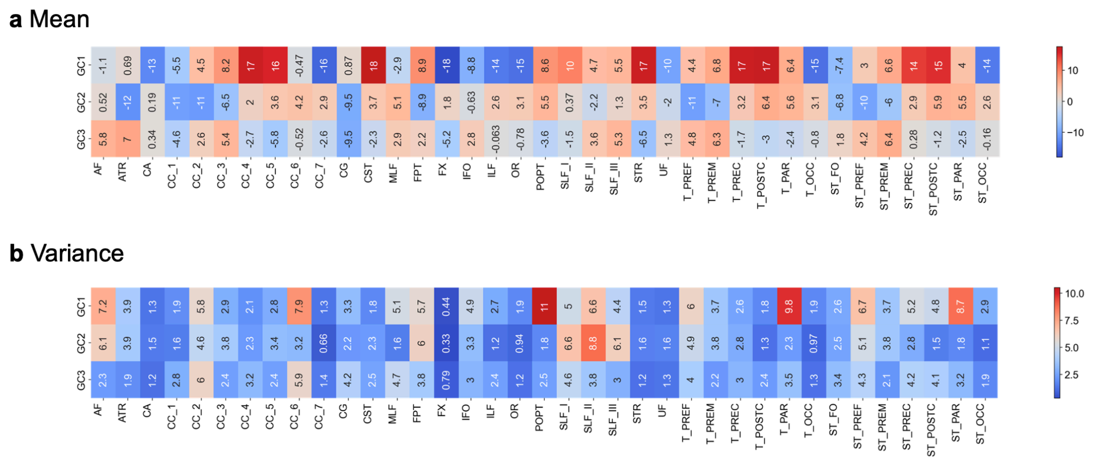


**Figure S7. Mean value and variance of fiber contributions to the global connectopies. (a)** The mean value of fiber contributions to the global connectopies. **(b)** Variance in fiber contributions to the global connectopies.

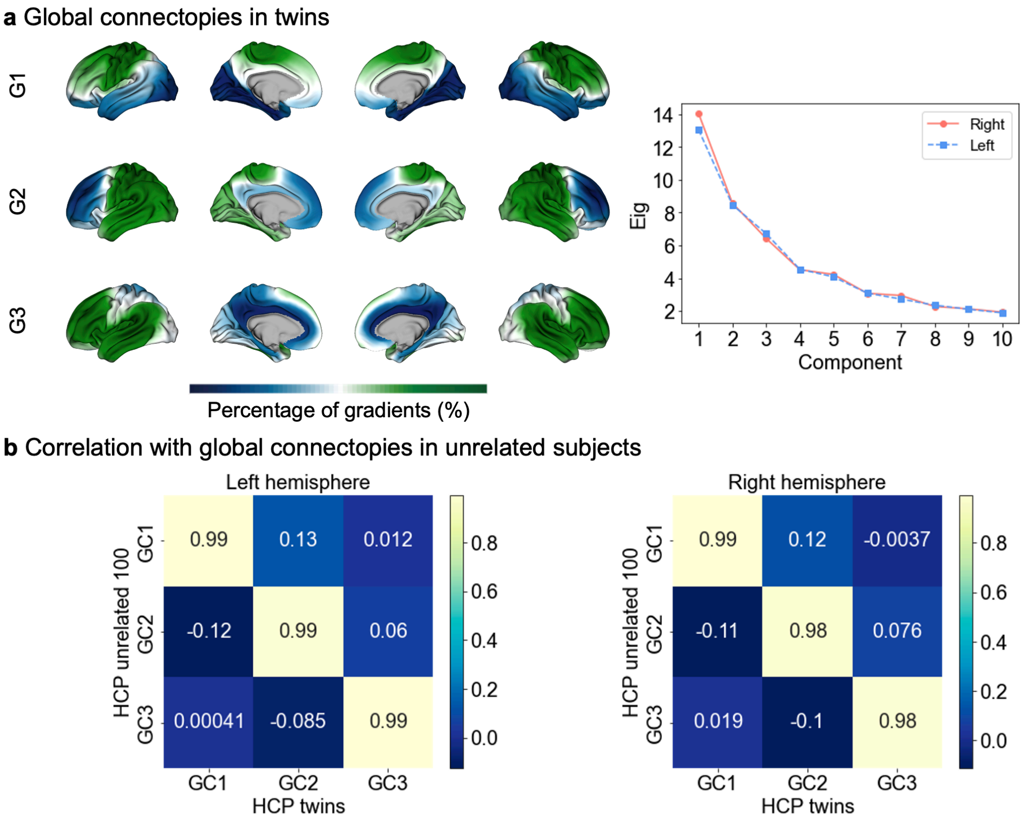


**Figure S8. Global connectopies were calculated using HCP twins and their correlation with global connectopies. (a)** We replicated our procedure of calculating global connectopies on 194 HCP twins. The scree plot describing variance is shown on the right. **(b)** The correlation between results on twins was very high with our original main results (left: $r_{G1\text{-}DV} = 0.99$, $r_{G2\text{-}RC} = 0.99$, $r_{G3\text{-}ML} = 0.99$; right: $r_{G1\text{-}DV} = 0.99$, $r_{G2\text{-}RC} = 0.98$, $r_{G3\text{-}ML} = 0.98$).

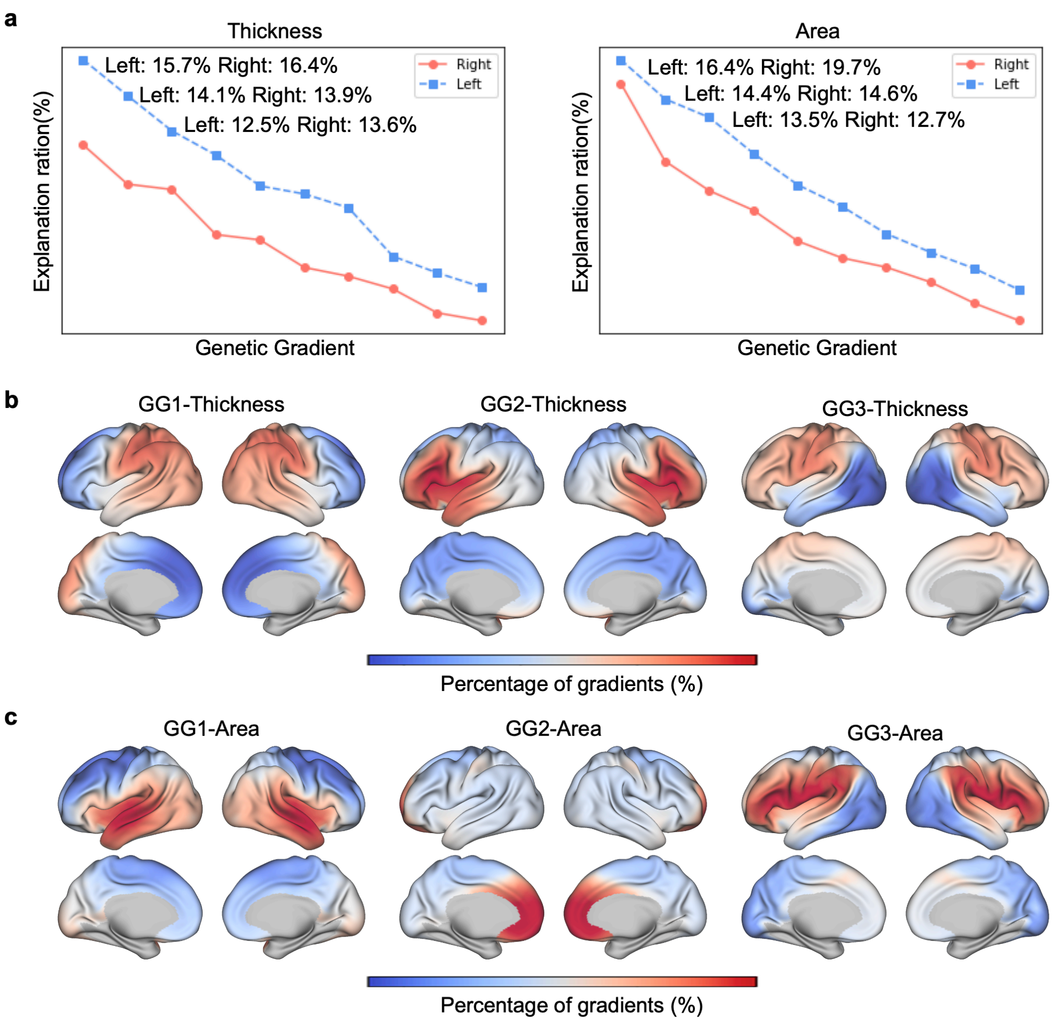


**Figure S9. Gradients of genetic similarity in cortical thickness and surface area of the cerebral cortex. (a)** Scree plot describing variance when calculating the gradients. **(b)** The first three gradients of genetic similarity of cortical thickness of the cerebral cortex. **(c)** The first three gradients of genetic similarity of the surface area of the cerebral cortex.

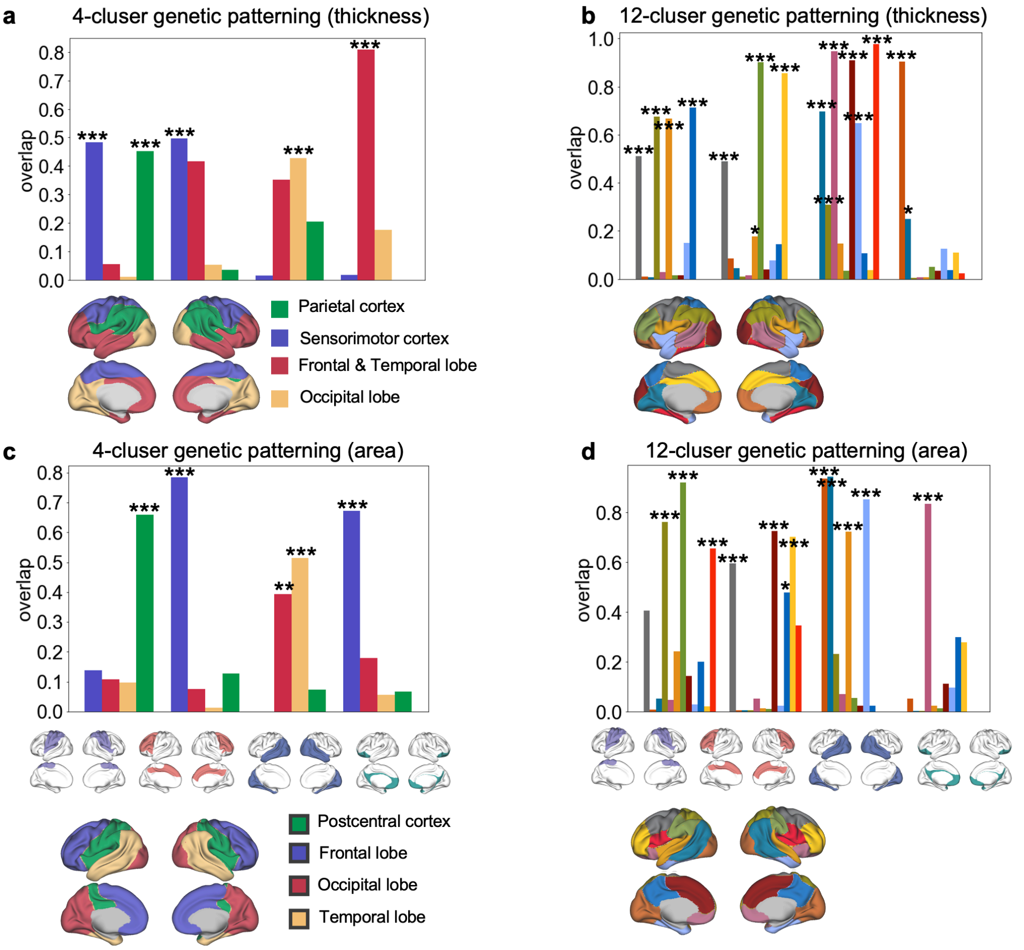


**Figure S10. Overlap between four modules derived from hierarchical clustering with genetic patterning.** Four modules derived from hierarchical clustering in Figure 3c in the main text showed significant overlap with one or more parcels in genetic patterning obtained from the genetic correlation of cortical thickness and surface area. *** $p < .001$, ** $p < .01$, * $p < .05$.

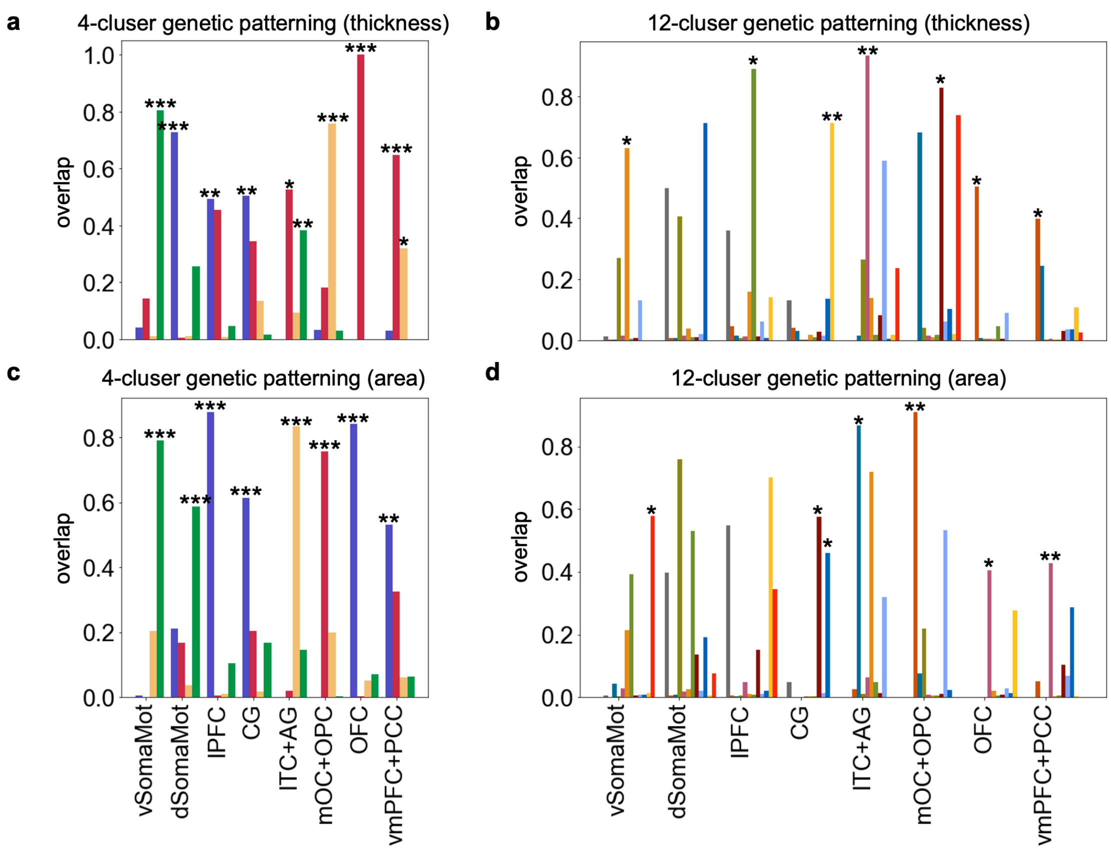


**Figure S11. Overlap between eight modules derived from hierarchical clustering with genetic patterning.** Eight modules derived from hierarchical clustering in Figure 3c in the main text showed significant overlap with one or more parcels in genetic patterning obtained from the genetic correlation of cortical thickness and surface area. *** $p < .001$, ** $p < .01$, * $p < .05$.

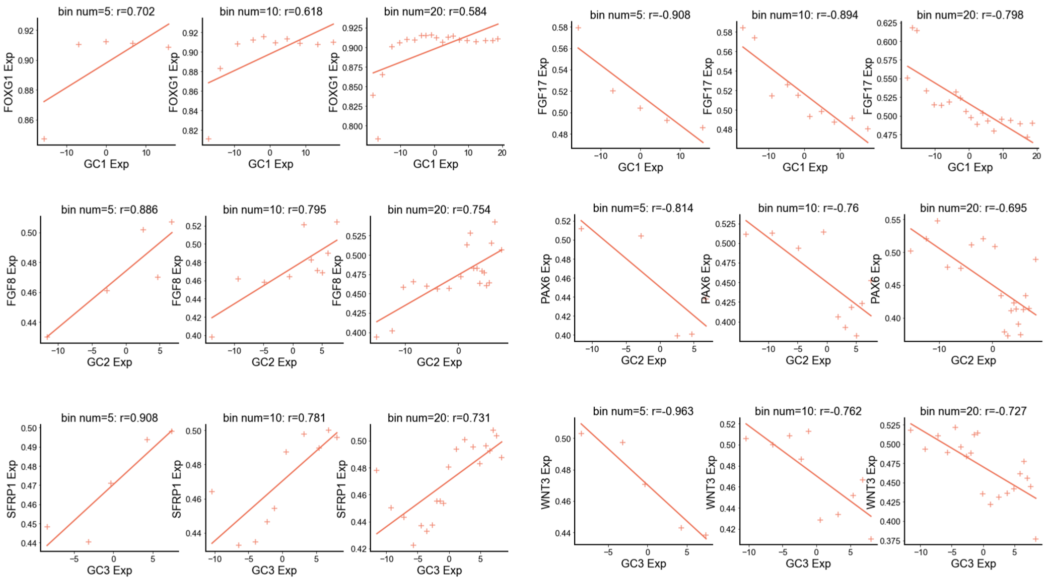

**Figure S12. Relationship between expression of example genes with the global connectopies.** We divided the connectopies and expressions of genes into bins (5, 10, and 20 bins) and calculated the correlation coefficient between these two. For GC1-DV, *FOXG1* and *FGF17* were selected. For GC2-RC, *FGF8* and *PAX6* were selected. For GC3-ML, *SFRP1* and *WNT3* were selected. We found a high correlation between connectopies and gene expression patterns.
